# Complement inhibition by C3-siRNA treatment prevents AChR loss and reduces complement activation in the rat Passive Transfer Myasthenia Gravis (PTMG)

**DOI:** 10.1101/2025.08.31.673367

**Authors:** Anja K Schöttler, Marina Mané-Damas, Britt Arets, Marc H. De Baets, Sandra MH Claessen, Tara Barbour, Erik Richardson, David Eyerman, Lukas Scheibler, Mario Losen, Pilar Martinez-Martinez

**Affiliations:** Department of Psychiatry and Neuropsychology, Mental Health and Neuroscience Research Institute, Maastricht University, the Netherlands; Apellis Pharmaceuticals, Inc., Waltham, Massachusetts, USA

**Keywords:** Myasthenia gravis, passive transfer, complement inhibition

## Abstract

Myasthenia gravis (MG), a well understood antibody-mediated autoimmune disease, is characterized by skeletal muscle weakness and fatigue. The autoantibodies are directed against neuromuscular junction (NMJ) proteins such as the acetylcholine receptor (AChR), essential for neuromuscular signal transmission. Anti-AChR antibodies are mainly of the immunoglobulin (Ig) types IgG1 and IgG3, which have a high capacity to activate the classical complement pathway. In AChR-MG, complement activation and the subsequent formation of the membrane attack complex (MAC) at the NMJ is one of the key effector mechanisms, causing substantial damage to the entire postsynaptic membrane and resulting in the typical clinical manifestations of muscle weakness and fatigue observed in both patients and animal models.

Since classical treatment strategies, which focus on general immunosuppression and enhancing the effect of ACh binding to the AChR, do not lead to full remission of symptoms in many cases, new treatment approaches are focusing more specifically on targeting effector mechanisms, such as complement activation. The first complement inhibitor approved for MG was eculizumab, a monoclonal antibody targeting C5. However, clinical trials showed that up to 28% of patients treated with eculizumab did not experience an improvement in myasthenic symptoms, emphasizing the urgent need for alternative therapies. Intervening earlier in the complement cascade might be more beneficial, as it could prevent the release of anaphylatoxins, such as C3a, that contribute to further inflammation at the NMJ.

In this study, the passive transfer myasthenia gravis (PTMG) rat model was used to investigate whether silencing hepatic C3 expression with a C3-targeted, small interfering (si)RNA, could ameliorate disease symptoms in the acute phase of MG. Female Lewis rats were injected subcutaneously with different dosages of C3-siRNA prior to PTMG induction with mAb35. C3-siRNA, administered weekly at a dose of 30 mg/kg, significantly prevented weight and AChR loss, and accordingly improved muscle function, as measured by muscle strength tests and electromyography. Consistently, the treatment reduced MAC deposition at the NMJ. Overall, these results provide insight into the efficacy of complement inhibitors in the acute phase of MG and suggest potential strategies for advanced treatment options in AChR-MG crisis.

## Introduction

Myasthenia gravis (MG) is an antibody-mediated autoimmune disease characterized by skeletal muscle weakness and fatigue. In MG, autoantibodies target proteins that are essential for neuromuscular signal transmission (1,2). In approximately 85% of MG cases, autoantibodies are directed against the acetylcholine receptor (AChR) at the postsynaptic membrane of the neuromuscular junction (NMJ)(3). AChR-MG is predominantly driven by immunoglobulins (Ig) of the IgG1 and IgG3 isotypes (4), which induce pathology through four major effector mechanisms: 1) Antigenic modulation, resulting in AChR internalization and a reduced receptor density due to autoantibody crosslinking (5); 2) Activation of the classical complement cascade and the formation of the membrane attack complex (MAC); 3) Blockage of the acetylcholine (ACh) binding site in the AChR (6–8), and 4) the loss of AChR associated proteins (9–12). While antigenic modulation and receptor blocking affect only the AChR, complement activation leads to significant damage to the entire postsynaptic membrane, resulting in the loss of both AChR and associated proteins (13,14).

Currently, first-line treatments for AChR-MG include acetylcholinesterase inhibitors (AChEI) such as pyridostigmine to counteract the degradation of ACh in the synaptic cleft, thereby enhancing signal transmission to the muscle (15). However, due to its short therapeutic effect of only 3-4 h, pyridostigmine is typically effective only in mild cases of MG (16). Patients often need additional immunosuppressive (IS) therapies such as corticosteroids (17). Unfortunately, side effects as well as low or non-responsiveness to IS therapy have been reported in 10-20% of MG patients (18). Furthermore, thymectomy is a routine procedure to reduce the required dose of immunotherapy in AChR-MG patients (17). While remission is frequently observed, surgical procedures carry inherent risks, and outcomes cannot always be reliably predicted (19).

Since the discovery of complement as a major pathogenic driver in MG, several experimental strategies have been approached to inhibit the classical complement pathway at various stages of the complement activation pathway, to prevent MAC deposition and consequential damage of the NMJ. These approaches include the use of synthetic peptides, monoclonal antibodies (mAb), and small interfering RNAs (siRNAs) (20–31). Commonly, for pre-clinical assessment of new MG therapies standardized MG rodent models are used. The passive transfer myasthenia gravis (PTMG) rat model is induced by injection of the monoclonal anti-AChR antibody mAb35 into the rats. This leads to muscle weakness, complement activation and ultrastructural changes of the NMJ (32). However, the PTMG model only addresses the acute effector phase of MG, without the lymphocytic response, while the experimental autoimmune myasthenia gravis (EAMG) rat and mouse model induced by active immunization of the animals mimic the full human disease progression (33,34). The administration of anti-C1q antibodies in the mouse EAMG model was effective at a dose of 10 µg, significantly reducing EAMG-related symptoms (35). Furthermore, EAMG mouse models deficient in complement factors C4 and C3 did not show complement deposition at the NMJ (36). Similar results were observed with siRNA targeting C2 which effectively reduced complement activation in EAMG in mice (25). Enhancing the activity of the decay-accelerating factor (DAF) to interrupt the classical pathway at the level of C3 and C5 convertases (37), as well as inhibiting MAC formation using C5- or C6-directed antibodies (21,26), C5-specific siRNA (27,38), or the C5-targeting protein rEV576 (24), led to reduced clinical symptoms and prevented MAC formation in PTMG rat model. Notably, Soltys *et al.* successfully replicated the beneficial effects of the treatment with rEV576 in the EAMG rat model (24). However, despite the beneficial effects in rodent models, to our knowledge only one C5-targeting siRNA advanced to clinical approval in human MG (39).

Eculizumab, a monoclonal antibody, and zilucoplan, a small synthetic peptide (31), both designed to block C5 cleavage, are approved therapies for the treatment of MG with AChR autoantibodies. Eculizumab is recommended for severe, refractory AChR-MG, as outlined in the 2020 MG management guidelines (29). Generally, targeting C5 and preserving upstream complement system functions would be beneficial to maintain the immune system functionality. However, targeting the C3 pathway in MG could provide broader impact by also preventing inflammatory processes mediated by the anaphylatoxin C3a, which is produced at the point of C3 cleavage/activation (42,43). The therapeutic potential of targeting C3 has been supported by several preclinical studies of MG, as well as in other autoimmune disease models. For example, in EAMG mouse models, C3-deficient mice showed reduced levels of anti-AChR antibodies in the blood and an absence of complement deposition at the NMJ (36). Similarly, in a mouse model of neuromyelitis optica, inhibition of C3 activation using a complement receptor 2 (CR2)-targeted CR2-complement receptor 1-related gene/protein y (Crry) fusion protein (CR2-Crry) restored motor functions and reduced inflammatory release (44). Additionally, Soltys *et al.* found that a deficiency of the C3 inhibitor, Crry, contributed to the development of murine EAMG (45), further demonstrating the relevance of C3 in AChR-MG pathogenesis. These findings underscore the potential of C3-targeting therapies to modulate disease mechanisms in AChR-MG and related autoimmune conditions.

Besides the choice of the target, the pharmacologic properties of the drug itself can also have an impact on the efficacy of the treatment. siRNAs are RNA duplexes, typically 17-27 base pairs in length, which induce degradation of a target messenger RNA (mRNA) via a mechanism known as RNA interference (RNAi) and thus reduce the target protein expression (46). RNAi is mediated through endogenous cellular machinery; the siRNA is loaded into a protein complex known as the RNA-induced silencing complex (RISC) and the siRNA guide strand binds to target mRNA through Watson-Crick base-pairing, allowing the endonucleolytic cleavage of the mRNA by the Ago2 protein in RISC. Due to this catalytic activity and slow release from endosomal compartments, siRNAs can exert therapeutic effects *in vivo* for up to 6 months (47), offering clear advantages for managing chronic disorders that currently require weekly or monthly interventions e.g. injections of the drug. Non-responders to eculizumab, for example, often exhibit genetic variations in C5, which reduce or negate the drug’s effectiveness (48). Developing new monoclonal antibodies is both time-consuming and costly, whereas the development of siRNAs is relatively cost-efficient and straightforward (49,50). Thus, siRNAs may expand opportunities to tailor treatments to specific genetic variants, potentially overcoming non-responsiveness to existing therapies. So far, siRNA studies targeting complement factors and regulatory proteins in experimental models of MG are encouraging (25,27,38).

Here, we studied the therapeutic efficacy of a siRNA targeting C3 (C3-siRNA) in the acute PTMG rat model. Hepatocytes are the primary site of synthesis of several circulating complement components, including C3. To achieve C3 suppression in liver, the siRNA was conjugated to a triantennary N-acetylgalactosamine (GalNAc) moiety, a ligand known to readily bind the asialoglycoprotein receptor (ASGPR), which is highly expressed in hepatocytes (51–53). We hypothesized that inhibiting complement activation by downregulating C3 production in the liver using GalNAc C3-siRNA would have a protective effect during the acute phase of MG and thereby prevent the development of characteristic myasthenic symptoms in the rat PTMG model.

## Materials and Methods

### C3-siRNA dose range finding study

A pilot study to determine the dose range of C3-siRNA to be administered in the PTMG study was performed by Apellis Pharmaceuticals in the Lovelace Biomedical’s animal research facilities. Twenty-four male Sprague Dawley rats (8-10 weeks old) were obtained from Charles River Laboratories, US, and injected subcutaneously (sc) with one single dose on day 1 or 3 daily doses on days 1, 2, and 3 of C3-siRNA, a nontargeting control siRNA, or PBS, as described in Table 1. Lyophilized siRNA was dissolved in PBS to concentrations of 0.2, 0.6, and 2 mg/ml to facilitate dose levels of 3, 10, and 30 mg/kg, respectively. Blood was collected in K2 EDTA tubes from the vena jugularis on day 0 (pre-dose), 3, 8, 15, 22, and 29 (termination). Blood was centrifuged for 10 min at 1300 g at 4 °C. Plasma was separated, immediately frozen and stored in aliquots at -80 °C. Plasma C3 levels were determined with a rat C3 ELISA Kit (Eagle Biosciences #RC321-K01) following the manufacturer instructions. In brief, 100 µl samples (diluted 1:10,000 in 1X dilution buffer) were incubated for 20 min at room temperature (RT) in the provided microtiter plate, aspirated, and washed four times with 1X wash solution. Subsequently, the samples were incubated for 20 min at RT in the dark with 100 µl of enzyme-antibody conjugate. After washing, 100 µl tetramethylbenzidine (TMB) substrate solution was added for 10 min at RT in the dark. The reaction was stopped with 100 µl stop solution, and the absorbance measured at 450 nm within 30 min.

**Table 1:** Experimental groups of the dose range study. Healthy Sprague Dawley rats were injected sc with C3-siRNA or placebo siRNA, dissolved in PBS respectively, or PBS. Blood plasma at different siRNA administration timepoints was collected and analysed to determine the circulating C3 levels.

| Number of animals | Treatment | Target Dose [mg/kg] | Cumulative dose [mg/kg] |
| --- | --- | --- | --- |
| 3 | C3-siRNA | 3 mg/kg | 3 mg/kg |
| 3 | C3-siRNA | 10 mg/kg | 10 mg/kg |
| 3 | C3-siRNA | 30 mg/kg | 30 mg/kg |
| 3 | C3-siRNA | 3x 1 mg/kg | 3 mg/kg |
| 3 | C3-siRNA | 3x 3.3 mg/kg | 10 mg/kg |
| 3 | C3-siRNA | 3x 10 mg/kg | 30 mg/kg |
| 3 | Placebo off-target siRNA | 3x 10 mg/kg | 30 mg/kg |
| 3 | PBS | 0 mg/kg | 0 mg/kg |

### PTMG study

This study included 48 female Lewis rats obtained from Charles River Laboratories, (Sulzfeld, Germany). All experiments were conducted with permission of the Animal Welfare Committee (CCD project license number: AVD107002017808), according to the Dutch laws, and in line with regulations of Maastricht University. At 9-weeks of age and before any intervention, animals were distributed into eight experimental groups (Table 2) with an equal group’s average weight.

**Table 2:** Experimental groups. Dissolved C3-siRNA at different doses or an equal volume of saline was injected on day 1 and day 8 sc, before PTMG induction on day 15. All animals were sacrificed on day 17.

| **PTMG / non-MG** | **Number of animals** | **Treatment; route** | **1<sup>st</sup> treatment Dose [mg/kg], day 1** | **2<sup>nd</sup> treatment Dose [mg/kg], day 8** |
| --- | --- | --- | --- | --- |
| Non-MG | 4 | C3-siRNA; sc | saline | 10 |
| Non-MG | 4 | C3-siRNA; sc | saline | 30 |
| Non-MG | 4 | C3-siRNA; sc | 30 | 30 |
| Non-MG | 4 | saline; sc | saline | saline |
| PTMG | 8 | C3-siRNA; sc | saline | 10 |
| PTMG | 8 | C3-siRNA; sc | saline | 30 |
| PTMG | 8 | C3-siRNA; sc | 30 | 30 |
| PTMG | 8 | saline; sc | saline | saline |

### Experimental design, treatment administration and PTMG induction

Baseline measurements for body weight, muscle strength (grip strength and hand climbing test) and blood sampling were performed after two weeks of acclimatization, at least one day before the first treatment injection, at the age of 9 weeks.

Lyophilized GalNAc C3-siRNA (Apellis Pharmaceuticals, MA, US) was dissolved in sterile RNase-free water to a final concentration of 13.3 µg/µl. Animals received either a single sc injection of i) 10 mg/kg, or ii) 30 mg/kg seven days before disease induction, on day 8, or iii) a total of two injections of 30 mg/kg, 7 and 14 days before disease induction, on day 1 and day 8, according to Table 2. Treatment control animals (saline) receiving no C3-siRNA were injected sc with 150 µl sterile 0.9% NaCl solution. PTMG was induced at 11-weeks of age by sc injection of 40 pmol/100 g body weight of mAb35 [deposited to the DSHB by Lindstrom, J. (Developmental Studies Hybridoma Bank (DSHB) product mAb35) on day 15, after C3-siRNA treatment. Rats were sacrificed on day 17, 48 hours after disease induction. Control animals (non-MG) were injected sc with an equal volume of saline (Fig. 2A).

### Weight and blood collection

Body weight was measured at baseline on day 0 and on days 4, 7, 11, 14, 15, 16, and 17. Additionally, animals were weighed before each treatment/saline injection to define the doses.

Blood was collected in K2 EDTA tubes (Sarstedt #16.444) from the vena saphena on days 0, 4, 7, 11, 14, and 17. Blood was processed within 30 min after venipuncture and centrifuged for 15 min at 900 g at RT. Plasma was separated, immediately frozen and stored in aliquots at -80 °C.

### Plasma C3 analysis

Plasma C3 levels were measured on days 0, 4, 7, 11, 14, and 17 by ELISA. Briefly, MULTI-ARRAY 96 Small Spot goat anti-rabbit IgG coated plates (Meso Scale Discovery (MSD) #L45RA) were blocked with 150 μl/well with 5% Blocker A buffer from (MSD #R93BA) for 1 h at RT with shaking (700 RPM). After washing 3 times with 300 μl/well washing buffer (PBS, 0.2% Tween), 25 μl/well of capture rabbit anti-rat C3 antibody (1 μg/ml, Hycult #HP8022-100UG) were added and incubated for 1-2 hours at RT with shaking or overnight at 4°C. Native rat C3 (CompTech #R113) was used as a standard for the assay, and both, standards and samples were diluted in 1% Blocker A buffer. Following a washing step as previously described, 25 μl/well standard or sample was added and incubated for 1 h at RT with shaking. Next, plates were again washed and 25 μl/well of primary detection goat anti-C3-Biotin antibody (1:1000 in 1% Blocker A buffer, Origene #AP21439BT-N) was added and incubated for 1 h at RT with shaking. After washing, 25 μl/well of sulfo-tagged streptavidin (1:1000 in 1% Blocker A buffer, MSD #R32AD-5) was added and incubated for 1 h at RT with shaking. After a final washing step, 150 μl/well of read Buffer T (diluted 1:4 in H_2_O, MSD #R92TC-1) was added and the plate was read using the MESO QuickPlex SQ 120 MM (MSD) according to manufacturer’s instruction. Discovery Workbench v4 software was used for data analysis and standard curve was plotted by 4-parameter curve fitting.

### Muscle strength measurement and disease scoring

Muscle strength and disease scoring were regularly assessed, on days 0, 4, 7, 11, and 14, after the blood collection and twice daily on days 15 - 17, after PTMG induction.

Briefly, all animals were assessed (while manually held near the base of the tail) for their capacity to grasp and hold the bar of the grip strength test meter GS3 (Panlab Harvard Apparatus, Barcelona, Spain), while they were retracted following the axis of the sensor. Each animal was retracted five consecutive times. Peak-force generation was recorded as pre-exercise muscle strength. Subsequently, animals were tested for their ability to climb onto the dorsal side of the experimenter’s hand while held by the base of the tail. Five attempts were allowed with a maximum of 1 min per attempt and 30 s resting intervals between attempts. The rat’s ability to climb onto the experimenter’s hand was scored as 0 - 3. Finally, five additional pulls on the grip strength meter were measured and documented as post-exercise muscle strength.

Clinical scoring was performed as described (32), based on the locomotion, general behavior and welfare: 0=no weakness; 1=fatigable or weakness observed only after exercise; 2=disease symptoms present before exercise, hunched posture, or head down; 3=no ability to grip, hindlimb paralysis, respiratory distress/apnea, or strong weight loss of 15% or more from the maximum body weight recorded, leading to humane endpoint (HEP). Animals were observed twice per test, and the average score per animal and timepoint calculated. For analysis, the average score per treatment group was calculated.

### Electromyography, termination of experiment, and organ collection

Analgesia with 0.05 mg/kg buprenorphine was injected sc 0.5–4 h before electromyography (EMG). Anesthesia was induced by inhalation of isoflurane mixed with oxygen and air in equal amounts. Isoflurane at 3-4% was used for induction and at 1.5-3% for maintenance. Electromyography was performed as previously described (10). In brief, the *nervus fibularis* (peroneal nerve) was stimulated with a series of 8 supramaximal stimuli of 3 Hz and a duration of 0.2 ms per stimulus. Compound muscle action potential (CMAP) measurements were repeated at intervals of 30 s. The recording was performed sc on the tibialis anterior (TA) muscle. Decrement was considered when the fourth CMAP of a recording showed a >10% decrease in amplitude (Fig. 5D, orange arrow) compared to the first CMAP in the same recording, for three consecutive recordings. The presence of decrement is directly correlated to the amount of functional AChR at the NMJ. During the procedure, the animal’s body temperature was kept between 36–38 °C using an infrared heating lamp (DISA, Copenhagen, Denmark) and a heating plate. Blood fractions (plasma), liver, diaphragm, both TAs, and brain cortex were collected after euthanasia.

### Quantitative immunofluorescence of NMJs

Quantitative immunofluorescence of NMJs was performed as previously described (9,10) with modifications. In brief, 10 µm isopentane-frozen TA muscle cryosections were stained for AChR (using α-bungarotoxin) and MAC or synaptic vesicle Protein 2 (SV2). Dried cryosections were fixed with 1% formaldehyde (Sigma Aldrich #F8775) in PBS for 10 min at RT, washed, and blocked with 5% *(v/v)* normal goat serum (NGS) in PBS for 30 min at RT. Slides were subsequently incubated overnight at 4 °C with: mouse anti-C5b-9 (4 µg/ml, Santa Cruz Biotechnologies #sc-66190) or mouse anti-SV2 (1.55 µg/ml, DSHB #AB_2315387). After washing, the slides were incubated for 1 h at RT with the biotinylated goat anti-mouse IgG (2.5 µg/ml, Jackson Immunoresearch #115-065-166). Finally, Streptavidin Alexa Fluor 594 (4 µg/ml, Invitrogen #S-32356), α-bungarotoxin Alexa Fluor 647 (3.33 µg/ml, Invitrogen #B35450) and Hoechst (1:1000, Sigma Aldrich #B2261) were applied for 1 h at RT. All antibodies were diluted in 2% *(w/v)* NGS in PBS and all washing steps followed a two-time incubation of 5 min in 0.05% *(v/v)* TritonX-100 in PBS and a third 5 min incubation in PBS, all at RT. Slides were mounted with 80% *(v/v)* glycerol in PBS.

The relative immunofluorescence intensities of AChR, SV2, and MAC at the NMJ were quantified as measures of their relative concentrations (9,10). Muscle sections from the TA of 6 PTMG-saline, 6 PTMG-2x 30mg/kg and 4 non-MG-saline animals were used (Table 2). In total, per animal four representative TA muscle sections from two different levels of depth of the TA, spaced 250 µm apart were analysed. Images were acquired with a fluorescence Olympus BX51WI microscope (Olympus, Hamburg, Germany), connected to a Hamamatsu EM-CCD C9100 camera (Hamamatsu Photonics K. K., Hamamatsu City, Japan), and controlled with Micromanager software (version 2.0.3) as previously described (54,55).

Endplate regions were identified based on their morphological structure and positive staining for AChR. Approximately 100 images per muscle (each containing 1 to 10 endplates) were acquired, and the fluorescence intensities for MAC, SV2 and AChR were quantified using ImageJ version 1.48a (NIH), in a blinded manner. For each muscle section, the mean intensities of AChR were normalized to SV2, and MAC intensities were normalized to AChR within the same area. The average of the normalized intensities from all endplates across a set of muscle sections was used for comparisons between treatment conditions.

### Qualitative immunofluorescence analysis of macrophage infiltration in NMJ

Immunofluorescence analysis for the macrophage marker CD68 along with AChR in TA muscles was performed. Ten µm isopentane-frozen TA muscle cryosections were stained for AChR (using α-bungarotoxin) and macrophages (CD68). Dried cryosections were fixed with 1% formaldehyde (Sigma Aldrich #F8775) in PBS for 10 min at RT, washed, and blocked with 5% *(v/v)* normal goat serum (NGS) in PBS for 30 min at RT. Slides were subsequently incubated overnight at 4 °C with mouse anti-CD68 (ED1) (2.5 µg/ml, Bio-Rad Laboratories # MCA341R). After washing, the slides were incubated for 1 h at RT with the biotinylated goat anti-mouse IgG (2.5 µg/ml, Jackson Immunoresearch #115-065-003). Finally, Streptavidin Alexa Fluor 594 (4 µg/ml, Invitrogen #S-32356), α-bungarotoxin Alexa Fluor 647 (3.33 µg/ml, Invitrogen #B35450) and Dapi (1:800) were applied for 1 h at RT. All antibodies were diluted in 2% *(w/v)* NGS in PBS and all washing steps followed a two-time incubation of 5 min in 0.05% *(v/v)* TritonX-100 in PBS and a third 5 min incubation in PBS, all at RT. Slides were mounted with 80% *(v/v)* glycerol in PBS.

### Radioimmunoassay for muscle AChR quantification

To quantify the AChR density in muscles, a radioimmunoassay (RIA) was performed, using ^125^I-Tyr54 labelled α-bungarotoxin (Perkin Elmer #NEX126H050UC). Snap frozen TA muscles were homogenized individually 3 times for 30 s with 30 s pause intervals on ice with an Ultra-Turrax homogenizer in 10 ml buffer A containing 0.01 M NaN_3_, 0.01 M EDTA, 0.2% iodoacetamide *(w/v)* and 1 mM phenylmethylsulfonyl fluoride (PMSF) in PBS. Homogenates were centrifuged at 22100 g for 30 min at 4 °C. The supernatants were disposed, and the pellets were resuspended in 1.25 ml of buffer B (buffer A + 2% *(v/v)* TritonX-100) for 45 s with the homogenizer. Homogenates were shaken on a reciprocal shaker at 280 RPM for 1 h at 4 °C, and subsequently centrifuged at 22100 g for 30 min at 4 °C. The supernatant was collected, aliquoted and stored at -80 °C until further use. For the RIA, 3 aliquots per muscle were used. To all tubes, 10 µl I^125^ α-bungarotoxin immunoprecipitation solution (1% ^125^I-α-bungarotoxin 50 µCi + 10% anti-AChR rat serum + 89% buffer B *(v/v)*) were added and incubated at 4 °C overnight. 100 µl goat anti-rat IgG polyclonal serum (Eurogentec, Belgium) was added and incubated for 4 h at 4 °C. The samples were washed 3 times with 1 ml washing buffer (0.5% *(v/v)* TritonX-100 in PBS) and subsequently centrifuged at 30000 g for 5 min at 4 °C. Gamma radiation was measured for 2 min per sample using the WIZARD^2^ 2480 Automatic Gamma Counter (Perkin Elmer).

### RT-qPCR for C3 mRNA expression in the muscle

C3 mRNA expression in the muscle was analyzed in a subset of animals. Considering the expectedly low intramuscular C3 expression, PTMG and non-MG animals of a second PTMG experiment (PTMG-2) with the project license number AVD10700202216263 were added to this analysis to enlarge the PTMG and non-MG control groups and counteract variability. The experimental conditions of the PTMG experiment are identical as described in the methods of this paper. In total, muscles of 10 non-MG animals, 20 PTMG animals, and 8 PTMG animals treated with 2 x 30 mg/kg C3-siRNA were included in this analysis.

RNA was extracted from 30 mg of rat TA muscle, then frozen and stored at -80 °C, using the RNeasy Fibrous Tissue Mini Kit (50) (Qiagen #74704) following the manufacturer’s instructions with adaptions. In brief, the tissue was homogenized in 2 ml tubes pre-filled with ceramic beads (Fisher Scientific #15-340-154) and 400 µl of Buffer RLT Plus containing 1% β-mercaptoethanol, with the Bead Ruptor ELITE (Omni #19-042E) run twice at 5 m/s for 1 min. Next, 590 µl RNase-free water and 10 µl Proteinase K solution provided with the kit were added to the lysate, mixed by pipetting, and incubated at 55 °C for 10 min before centrifugation. The supernatant was transferred to a new microcentrifuge tube and 0.5x lysate-volume of 96-100% ethanol was added and mixed by pipetting for lysate clearance. A volume of 700 µl lysate was centrifuged in a RNeasy Mini spin column until all lysate was processed. The column membrane was washed once by centrifugation with 350 µl Buffer RW1, the flow through was discarded. To denature DNA, 10 µl DNase 1 stock solution in 70 µl Buffer RDD were added to the sample and incubated for 15 min at RT, eventually the reaction was stopped with 350 µl Buffer RW1 and centrifuged. After washing the sample twice with 500 µl Buffer RPE (with prolonged centrifugation time to 2 min the second wash), RNA was eluted in 40 µl RNase-free water by centrifugation for 1 min. All centrifugation steps were performed at RT for 15 s at >8000 g, if not indicated differently. All steps were performed in RNase-free conditions. The RNA concentration was determined using Nanodrop. Absorption ratios of A260/A280 >1.9 were considered appropriate for RT-qPCR analysis. RNA was stored for further processing at -20 °C.

The reverse transcription was performed using the SuperScript™ IV VILO™ Master Mix (Invitrogen #11766050), following the manufacturers instructions. Briefly, each sample’s cDNA digestion mix was prepared with 1000 ng RNA, 1 µl 10x ezDNase buffer, 1 µl ezDNase, and RNase-free water up to 10 µl, incubated for 2 min at 37 °C and stored on ice. 4 µl SuperScript™ IV VILO™ Master Mix and 6 µl RNase-free water were added to the reaction tube on ice, mixed, and incubated for 10 min at 25°C to anneal the primers, followed by the reverse transcription to cDNA for 10 min at 50 °C and the final enzyme activation for 5 min at 85 °C. The final cDNA was stored at -70 °C until further processing.

For the RT-qPCR analysis of C3 gene expression in the muscles, the TaqMan™ Multiplex Master Mix (Applied Biosystems #4461881) was used, following the manufacturer’s instructions of the TaqMan™ Fast Advanced Master Mix (Applied Biosystems #4444964). In brief, 2 µl cDNA 1:10 diluted in RNase-free water, 5 µl of 2x TaqMan™ Multiplex Master Mix, and 0.5 µl of each primer C3 assay (FAM Rn00566466_m1) and ActB assay (VIC Rn00667869_m1) were gently mixed and toped up to a total volume of 10 µl with nuclease-free water. After short vortex and spin down, the RT-qPCR was run in the CFX Opus 384 Real-Time PCR System (BioRad #12011452): polymerase activation at 95 °C for 20 sec, followed by 40 cycles of denaturation for 1 sec at 95 °C plus annealing/extension for 20 sec at 60 °C. The results were analysed with the delta-delta Ct method.

### Statistical analysis

GraphPad Prism 10 was used for statistical analysis. A Two-Way Analysis of Variance (ANOVA) was applied to individual treatment groups, followed by a One-Way ANOVA for normally distributed values, with Bonferroni’s multiple comparison as the post hoc analysis. Clinical scores were analyzed using the Kruskal-Wallis test with multiple comparisons. A two-sided p-value of 0.05 or less was considered statistically significant.

## Results

### C3-siRNA Dose Finding Study

Prior to the PTMG experiment, different doses of C3-siRNA were tested in healthy Sprague Dawley rats to establish the dose range for the PTMG study. As expected, PBS-treated animals did not show any reduction in plasma C3 levels throughout the pilot study. In contrast, single injections of C3-siRNA demonstrated a clear dose-dependent reduction in plasma C3 levels as early as day 3 (Fig. 1). The lowest C3 levels were detected on day 8, with plasma C3 reduced to 43%, 9% and <2% of baseline C3 levels following single injections of 3, 10, or 30 mg/kg C3-siRNA, respectively. Notably, in animals receiving the 30 mg/kg C3-siRNA dose, C3 levels remained stable at <2% until day 15, before gradually increasing. In contrast, animals treated with lower doses (3 and 10 mg/kg), showed a rise in C3 levels beginning after day 8. Cumulative doses of 3, 10 and 30 mg/kg C3-siRNA, administered as three consecutive injections on days 1, 2 and 3, yielded results comparable to those observed with single injections (Supplementary Fig. S1). In summary, doses of 10 and 30 mg/kg C3-siRNA both clearly reduced C3 protein levels in plasma and were included in the PTMG C3-siRNA study.

**Figure 1:**
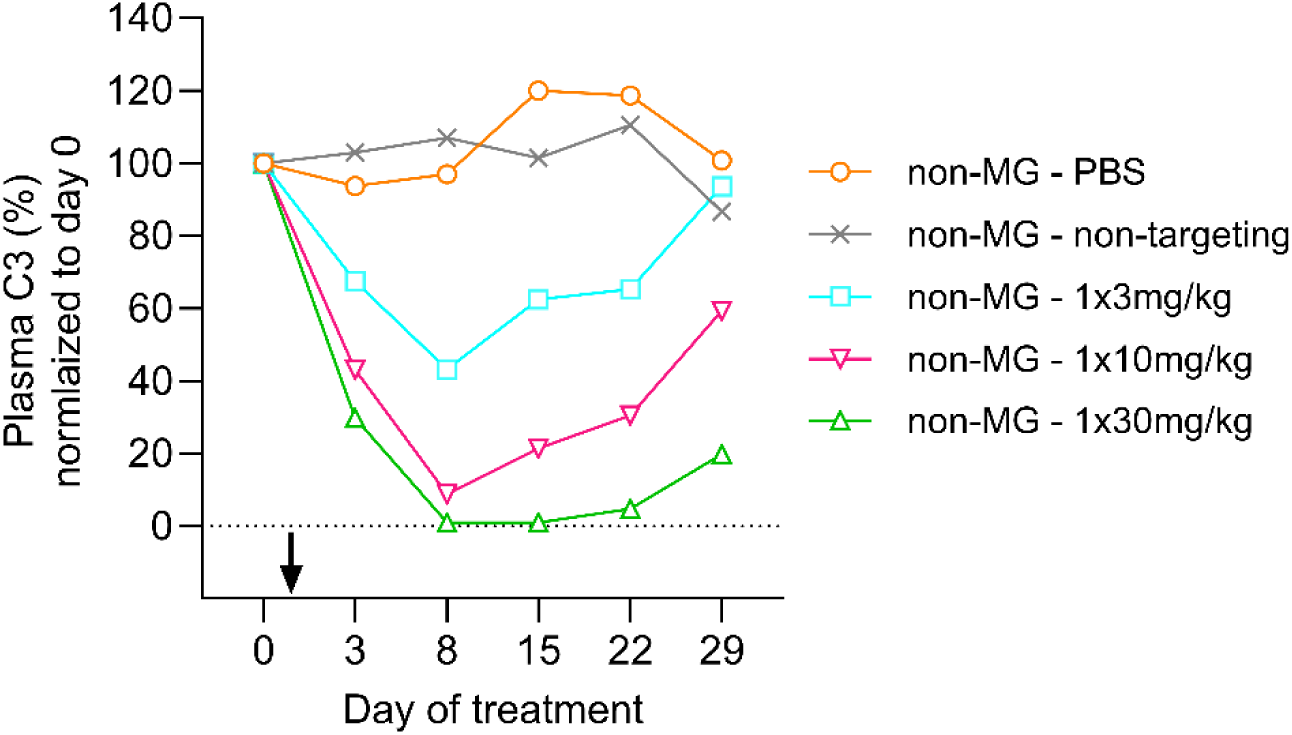
**C3 protein levels in plasma** of healthy animals receiving a single treatment injection to establish the dose range of C3-siRNA for the PTMG study. After baseline measurements on day 0, C3-siRNA was administered on day 1 (black arrow) and plasma C3 protein levels analyzed on days 3, 8, 15, 22, and 29. Per group N=3. Group averages are shown.

Additional analyses confirmed that hepatic C3 mRNA expression was effectively inhibited by the C3-siRNA (Supplementary Fig. 5)

### C3-siRNA Elicits a Dose-Dependent Reduction in Circulating C3 Levels

Circulating C3 protein levels were measured in plasma samples, using an MSD-based ELISA method (Fig. 2). C3 levels were decreased consistently in a dose-dependent manner, by 4 days post-dose and remained low throughout the study. Animals treated with the 2x30 mg/kg dose showed the largest decrease, as expected (Fig. 2A-D). C3-siRNA was detected in the liver of PTMG animals that received a single 30 mg/kg C3-siRNA injection, while no C3-siRNA was detected in non-MG saline-treated animals, as expected. One PTMG animal treated with a single 30 mg/kg C3-siRNA injection did not exhibit any reduction in plasma C3, suggesting a missed injection. Consequently, this animal was excluded from the analysis, resulting in a total of N=7 animals for that treatment group. In the excluded animal, no C3-siRNA was detected in the liver resulting in C3 levels similar to those of saline injected animals (Supplementary Fig. S2).

**Figure 2:**
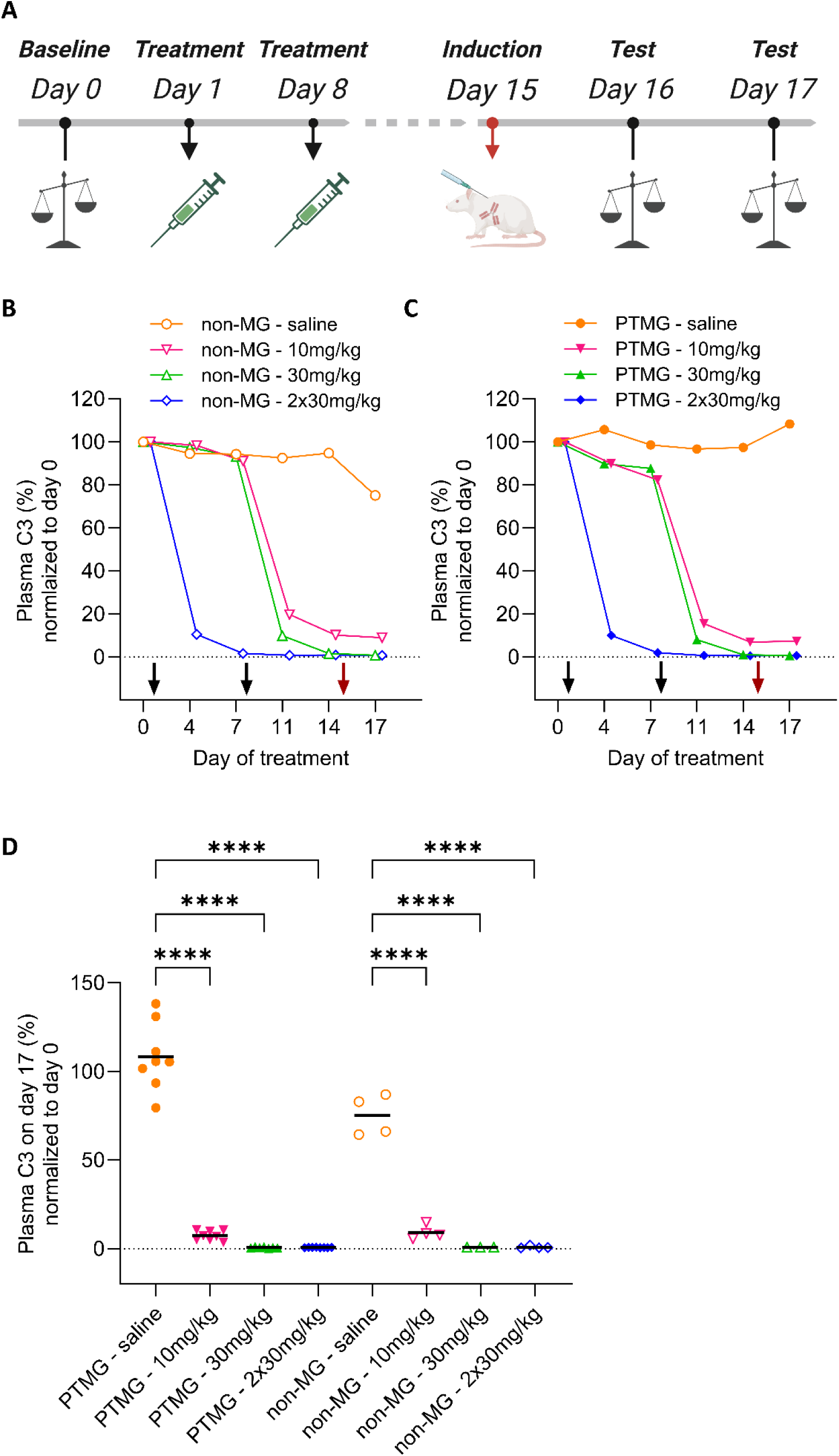
General experimental design and C3 protein levels in plasma. **A)** After baseline measurements on day 0, subcutaneous treatment injections were administered on day 1 and 8. Intermediate testing and blood collection were performed on day 4, 7 and 11 (not shown). On day 15, indicated with a red arrow, PTMG was induced using mAb35, while non-MG animals received a saline injection of the same volume. **B-C)** Plasma C3 expression was reduced by the sc administration of C3-siRNA at various dose levels in **B)** non-MG animals and **C)** PTMG animals. For animals receiving 2 x 30 mg/kg treatment, doses were administered on days 1 and 8. For animals receiving 1 x 10 or 1 x 30 mg/kg doses, administration occurred on day 8, as indicated by the black arrows on the graphs. Saline-treated animals showed no significant change in C3 levels over time. Group averages are shown. **D)** Plasma C3 level changes in individual treatment groups on day 17, normalized to baseline levels on day 0. Individual plasma C3 levels per animal are shown. Statistical analysis was performed with Two-Way ANOVA (B-D), followed by One-Way ANOVA and post-hoc Bonferroni’s multiple comparison test (D); α=0.05; ****p<0.0001. A) Created in BioRender. Schöttler, A. (2025) https://BioRender.com/b4xxodd.

### C3-siRNA Prevented Bodyweight Loss in PTMG animals

Bodyweight was recorded at least twice per week throughout the study. All animals gained weight similarly until PTMG induction on day 15, regardless of the treatment group (Fig. 3). Treatment with C3-siRNA did not affect weight progression in non-MG animals at any dose level or time point. Following PTMG induction or saline injection for non-MG animals, bodyweight plateaued in non-MG animals (Fig. 3A). In PTMG animals, weight either plateaued or decreased depending on the treatment received (Fig. 3B). Bodyweight loss was significantly prevented in PTMG animals treated with C3-siRNA compared to saline-treated PTMG animals, showing a clear dose dependent effect (Fig. 3B). Before euthanasia on day 17, no loss in bodyweight was observed in animals treated with the double dose of 30 mg/kg C3-siRNA (p<0.0001), while animals receiving a single injection of 30 mg/kg C3-siRNA presented a reduced weight loss of only 2% (p=<0.0001). Animals treated with the low dose of 10 mg/kg C3-siRNA exhibited a 6% weight loss (p=0.0441), whereas saline-treated PTMG rats experienced substantial weight loss of approximately 10% (Fig. 3C).

**Figure 3:**
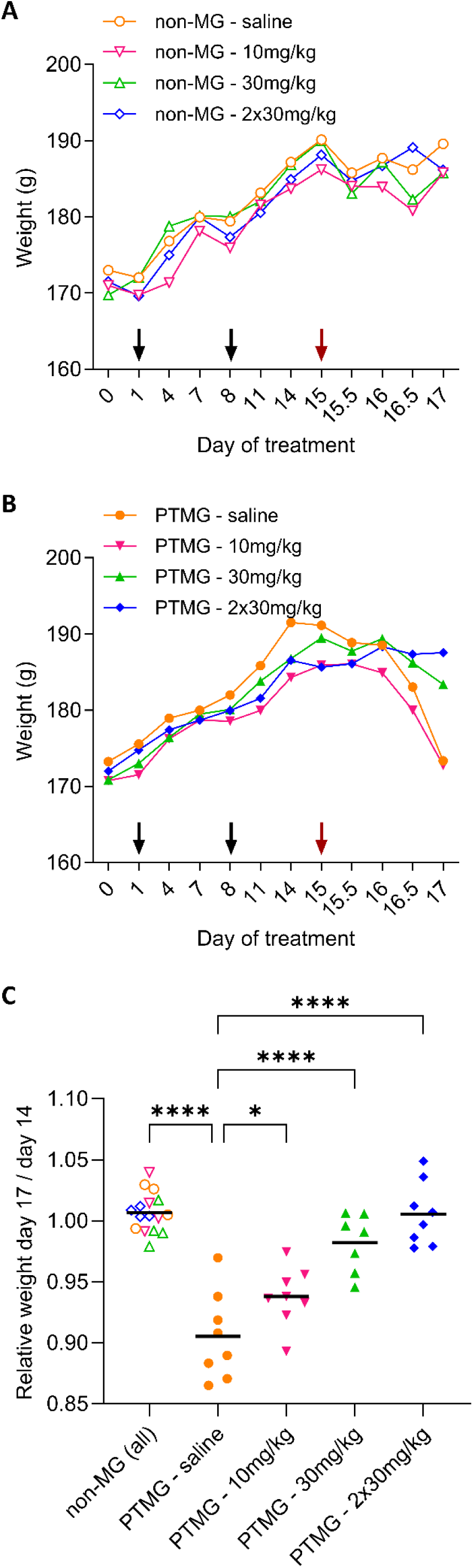
Weight progression during the experiment. Weight development of **A)** non-MG and **B)** PTMG animals, showing a clear dose-dependent effect in the PTMG groups. **C)** The relative weight on day 17 compared to day 14 was calculated, with all non-MG animals pooled into one group. Each dot represents an individual animal. Statistical analysis was performed using Two-Way ANOVA (A-B), followed by One-Way ANOVA and post-hoc Bonferroni’s multiple comparison test; α=0.05; *p=0.0441, ****p<0.0001.

### C3-siRNA Reduced Disease-Specific Muscle Weakness/Clinical Manifestations in PTMG animals

Muscle strength was measured by assessing grip strength, and disease severity was classified based on muscle weakness in all animals. First, we observed an increase in grip strength after baseline measurements in all animals, which we attributed to training and muscle maturation (Fig. 4A). As expected, following PTMG induction on day 15, grip strength significantly differed between the treatment groups (Fig. 4 A-B). Non-MG animals exhibited a strong grip of approximately 9 Newtons (N) before and after disease induction. In contrast, saline-treated PTMG animals experience a loss of about 55% in grip strength on day 17, decreasing from an average of 9.6 N to 4.3 N. PTMG animals treated with 10 mg/kg C3-siRNA showed a 41% loss of grip strength on day 17, which was not significantly different from saline-treated PTMG animals. However, PTMG animals treated with single and double doses of 30 mg/kg C3-siRNA retained more muscle strength, losing only 12% and 14% of their grip strength, respectively, with no significant difference between the two groups. Both single and double 30 mg/kg C3-siRNA groups were significantly different from the saline-treated PTMG animals (Fig. 4B, p<0.0001).

**Figure 4:**
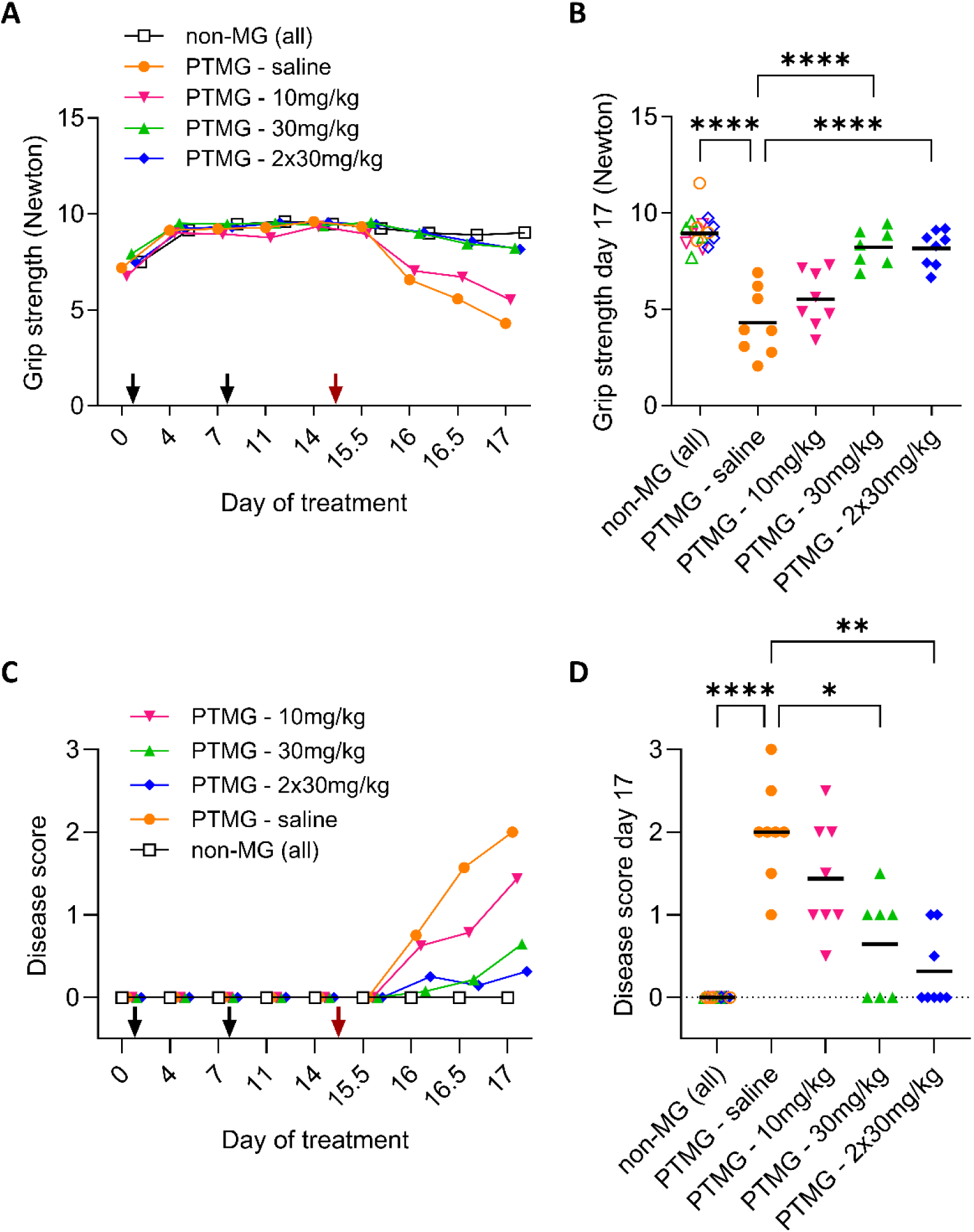
Progression of muscle weakness indicated by grip strength in the front limbs and subsequent disease scoring. **A)** Grip strength decreased in PTMG animals after disease induction (red arrow) with a dose-dependent effect of the C3-siRNA treatment. **B)** Grip strength on day 17 with non-MG animals pooled into one group. **C)** Disease scoring increased in PTMG animals after disease induction (red arrow), with an evident dose dependent effect of the C3-siRNA treatment. **D)** Disease score on day 17 with non-MG animals pooled into one group. In A) and C) group averages are shown. In B) and D) each dot represents one individual animal; horizontal bars indicate group means. Statistical analysis was performed with Two-Way ANOVA (A-C), followed by One-Way ANOVA and post-hoc Bonferroni’s multiple comparison test (B) or Kruskal-Wallis test (D); α=0.05; *p=0.0408, **p=0.0021, ****p<0.0001.

Correspondingly, PTMG animals showed clear differences in their disease scores depending on the treatment (Fig. 4C-D). The onset of muscle weakness occurred on day 16, one day after disease induction, in both the saline and 10 mg/kg C3-siRNA groups and progressing to severe muscle weakness before exercise. These animals reached, on average, disease scores of 2 and 1, respectively, by day 17 before euthanasia. In contrast, animals treated with single and double 30 mg/kg C3-siRNA doses showed an improvement in disease score compared to the low dose or saline treated PTMG animals. On day 17, muscle weakness was prevented in these groups with weakness only manifesting after exercise (average score = 0.6) or with no weakness (average score = 0.3) in each group, respectively (Fig. 4D).

### C3-siRNA Reduced Electrophysiological Abnormalities Associated with MG

EMG was performed before euthanasia, with decrement serving as an indirect measure of functional muscle AChR content. Both single and double 30 mg/kg C3-siRNA treatments significantly reduced the decrement to 6% and 4% respectively, compared to saline-treated PTMG animals (both p<0.0001). Consistent with the previously reported clinical manifestations, animals treated with 10 mg/kg C3-siRNA and saline-treated PTMG animals showed a strong decrement of 17% and 21% respectively, which were significantly different from non-MG animals (Fig. 5A). Non-MG animals showed, as expected, a slight increase of 2%.

**Figure 5:**
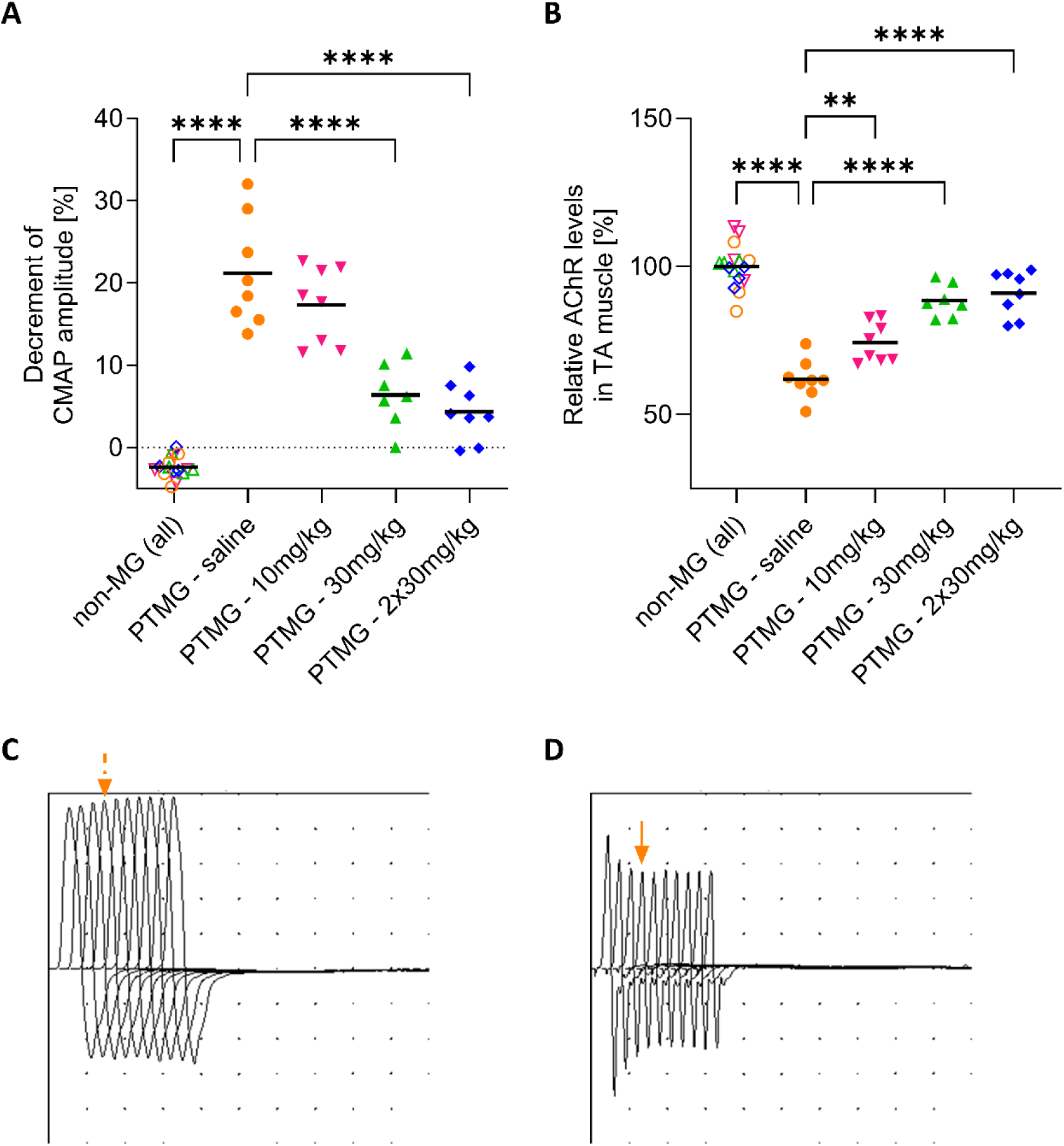
Disruption of the NMJ indicated by decrement recorded during EMG and AChR levels in the TA muscle by RIA. **A)** EMG was performed during terminal experiments with 3 Hz stimuli. Decrement was calculated as the reduction in amplitude of the fourth CMAP compared to the first CMAP of the same recording. The average of three consecutive recordings per animal was calculated and represented by a dot. **B)** Muscle membrane extracts containing AChR were incubated with ^125^I-labelled α-bungarotoxin, and radioactivity was measured in counts per minute (CPM). **C–D)** Representative images of CMAP recordings in **C)** healthy rats with slight increment of the fourth CMAP (dashed, orange arrow), and **D)** PTMG rats with ca. 25% decrement of the fourth CMAP (continuous, orange arrow). Each dot represents CPM per animal, normalized to the average CPM of all pooled non-MG animals. A Two-Way ANOVA for all individual groups was performed, followed by a One-Way ANOVA with post-hoc Bonferroni’s multiple comparison test; α=0.05; **p=0.0037, ****p<0.0001.

### C3-siRNA Reduced Muscle AChR Loss In PTMG Rats

Rat tibialis anterior muscles were used to quantify muscle AChR content, as its reduction is a hallmark of MG (Fig. 5B). AChR levels in TA muscles of saline-treated PTMG rats were reduced by nearly 38% compared to non-MG control animals (p<0.0001). In contrast, muscle tissue from rats receiving a single or double injection of 30 mg/kg C3-siRNA had 26% and 29% higher AChR levels, respectively, compared to saline-treated PTMG animals (both p<0.0001). Consistent with previous results, a single injection of 10 mg/kg C3-siRNA was less effective but still significantly prevented AChR loss compared to saline-treated PTMG animals (p=0.0004).

### Reduction in Complement Deposition at the NMJ in C3-siRNA Treated PTMG Rats

Quantitative immunofluorescence was used to evaluate the content of AChR, MAC, and SV2 at the NMJ. Six animals treated with a double injection of 30 mg/kg C3-siRNA, six animals of PTMG-saline and four non-MG saline-treated animals were included in the assessment.

The presynaptic protein, SV2, which remains unaffected by MG or PTMG (Supplementary. Fig. S3), was used for normalization and to assess AChR intensities. Non-MG animals exhibited strong AChR intensity, considered as 100%, representing the AChR content of healthy animals. Induction of PTMG in saline-treated animals significantly reduced AChR content at the NMJ (p<0.0001), with a reduction of 62% of the AChR content compared to non-MG animals. Treatment with two injections of 30 mg/kg C3-siRNA significantly protected from PTMG induced AChR loss. PTMG animals treated with two 30 mg/kg injections of C3-siRNA exhibited 68% AChR fluorescence intensity compared to healthy controls, significantly higher than that of saline-treated PTMG animals (p=0.0093) (Fig. 6B, C). However, it did not completely mitigate AChR loss, since treated muscles presented with 32% reduction in AChR levels compared to non-MG animals.

**Figure 6:**
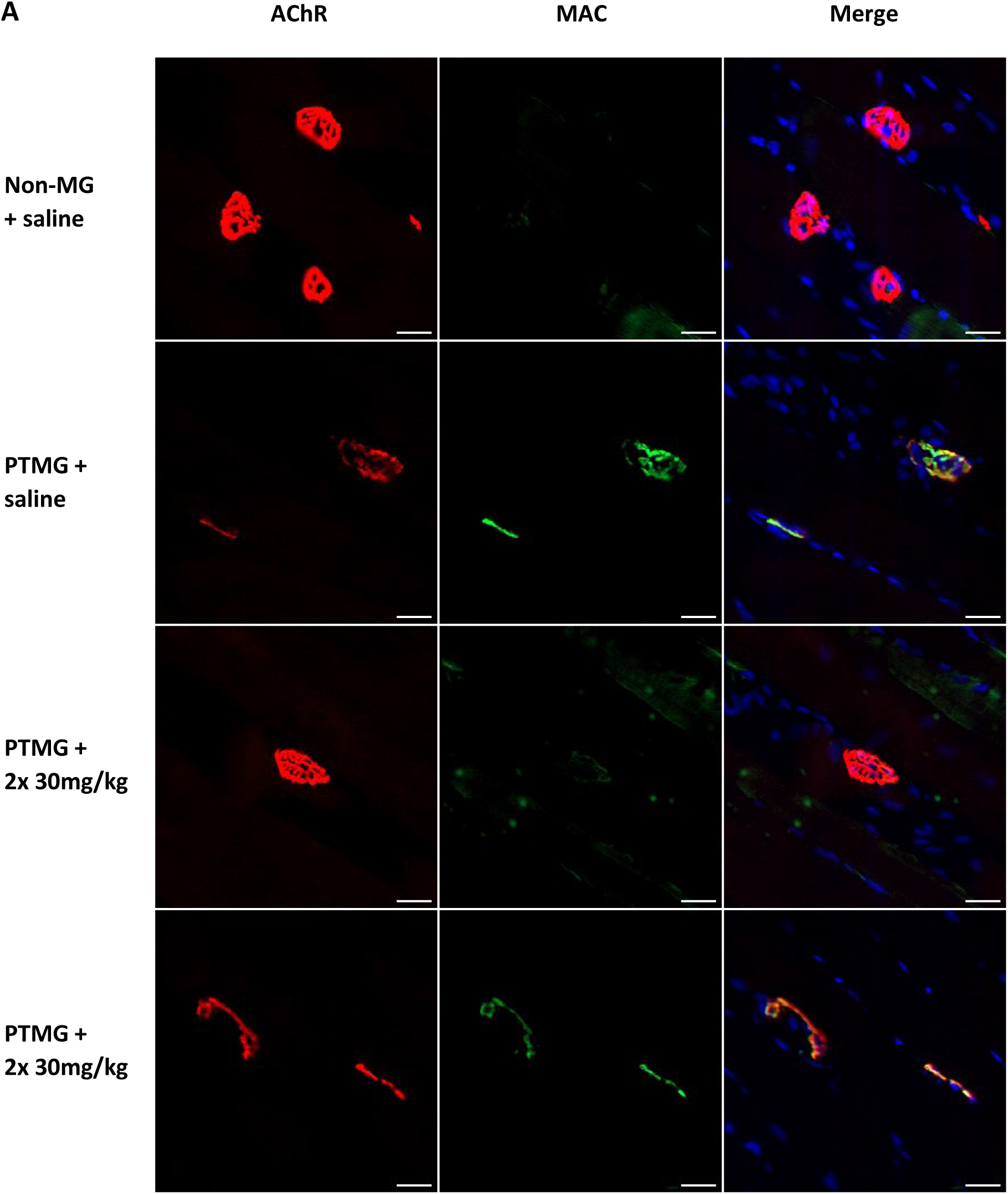
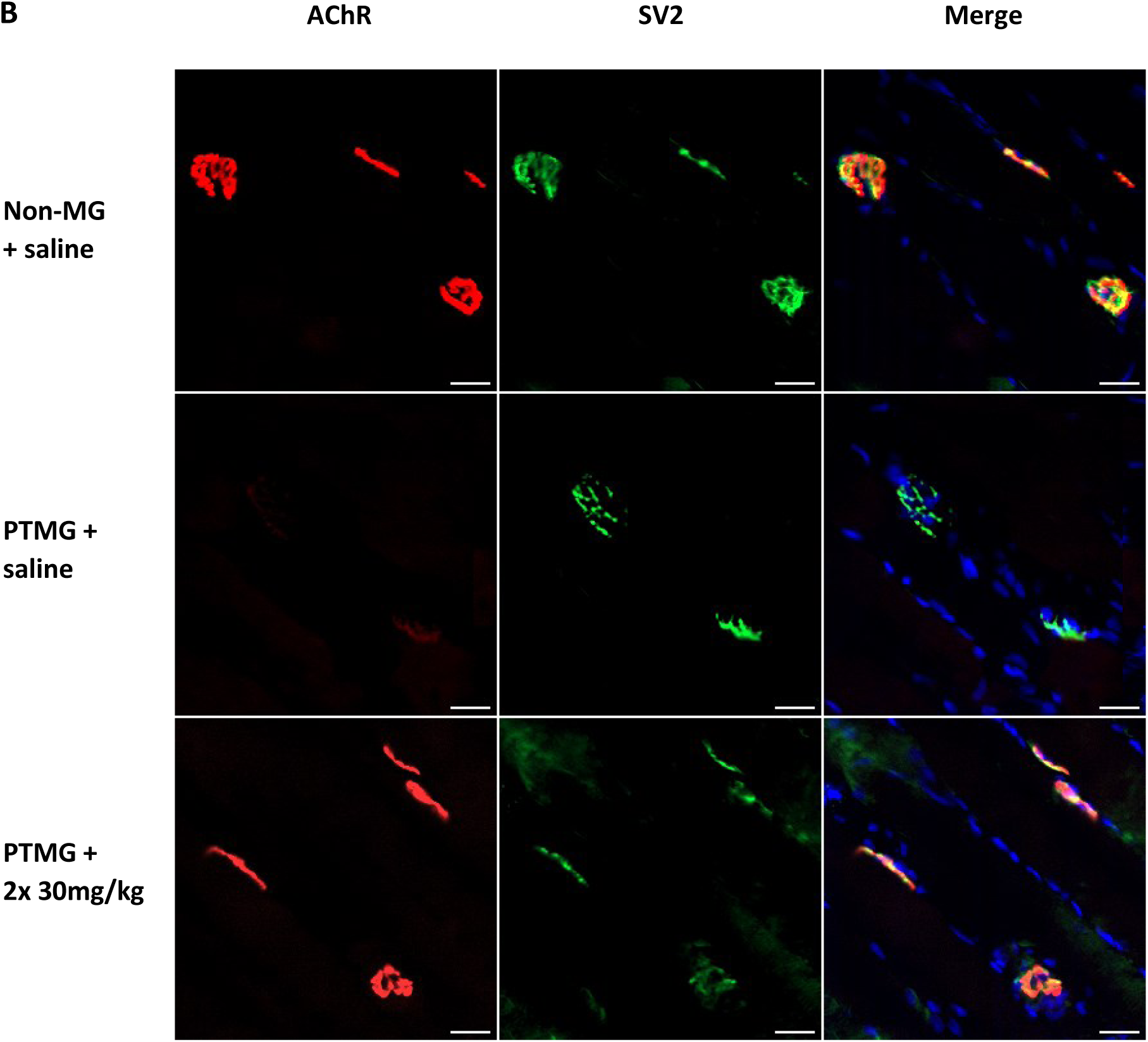
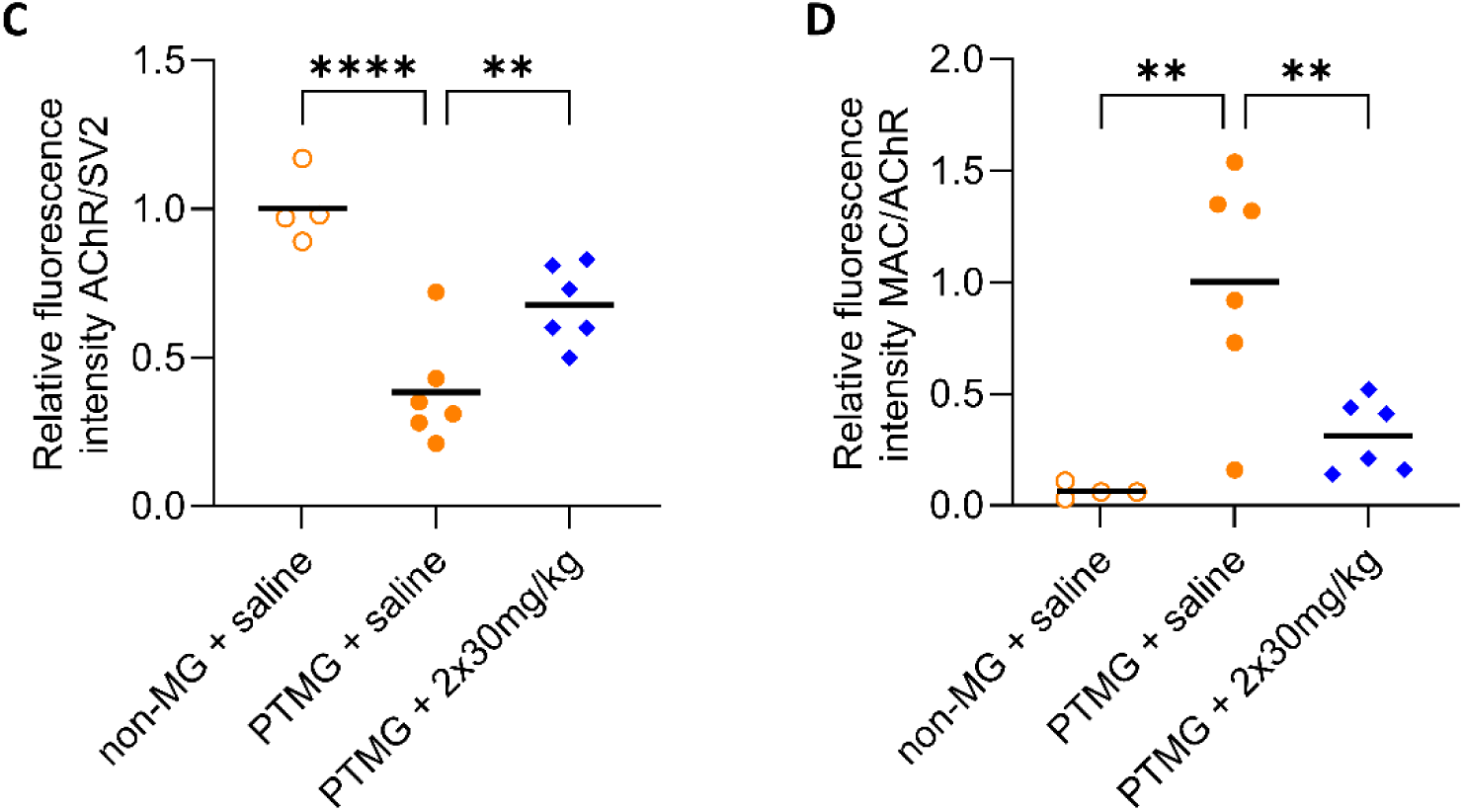
Fluorescence Microscopy Images of Representative Endplates and Quantitative Immunofluorescence Analysis. **A-B)** Representative NMJ images from the non-MG-saline, PTMG-saline, and PTMG 2x 30mg/kg treated groups. Muscle sections were stained for Hoechst (blue), AChR (red) and either **A)** MAC or **B)** SV2 (green). **C-D)** Quantification of relative fluorescence intensity of **C)** AChR expression normalized for SV2 and **D)** MAC levels normalized to AChR. Groups in C were normalized to the non-MG, saline treated animals. Groups in D were normalized to the PTMG, saline treated animals. N=4 non-MG animals, N=6 PTMG-saline animals, N=6 PTMG-2x 30 mg/kg animals. Each dot indicates one animal. Statistical analysis was performed using One-Way ANOVA with post-hoc Bonferroni’s multiple comparison test; α=0.05; **p<0.0094, ****p<0.0001. Scale bar = 25 µm.

AChR and MAC were visualized in the same NMJs (Fig. 6A, D). No MAC deposition at the NMJ was detected in non-MG animals. As expected, the highest MAC deposition was observed in saline treated PTMG animals, which was considered as 100% fluorescence intensity. Treatment with two 30 mg/kg injections of C3-siRNA significantly reduced MAC deposition, with a 31% decrease in fluorescence intensity of levels observed compared to the saline treated PTMG animals (p=0.0065). The remaining MAC deposition correlates with the AChR loss in the animals receiving two injections of 30 mg/kg C3-siRNA and indicates that while C3-siRNA treatment reduced MAC deposition at the NMJ, it did not prevent MAC deposition entirely.

### Muscle C3 mRNA expression can be reduced with C3-siRNA treatment in PTMG animals

C3 gene expression levels in muscle tissue from non-MG animals were considered as naïve expression and used for data normalization. PTMG-saline treated animals expressed significantly more C3 mRNA in TA muscles compared to non-MG animals (p<0.0001). PTMG animals treated with a double dose of 30 mg/kg C3-siRNA expressed significantly reduced C3 mRNA in their TA muscles compared to PTMG-saline treated animals, however, C3 expression in these animals is higher than in non-MG animals, even though not statistically relevant, likely due to high variability.

### PTMG related macrophage infiltration in muscles is not prevented by C3-siRNA treatment

Qualitative immunofluorescence analysis for the macrophage marker CD68 along with AChR was performed in TA muscles to assess the capacity of the C3-siRNA treatment to prevent PTMG-induced macrophage infiltration into muscles. As expected, no macrophages were found in non-MG animals. In contrast, both C3-siRNA and saline-treated PTMG animals presented with intense macrophage infiltration near the NMJ (Fig. 8), indicating that C3-siRNA treatment did not prevent macrophage activation in response to PTMG induction.

**Figure 7:**
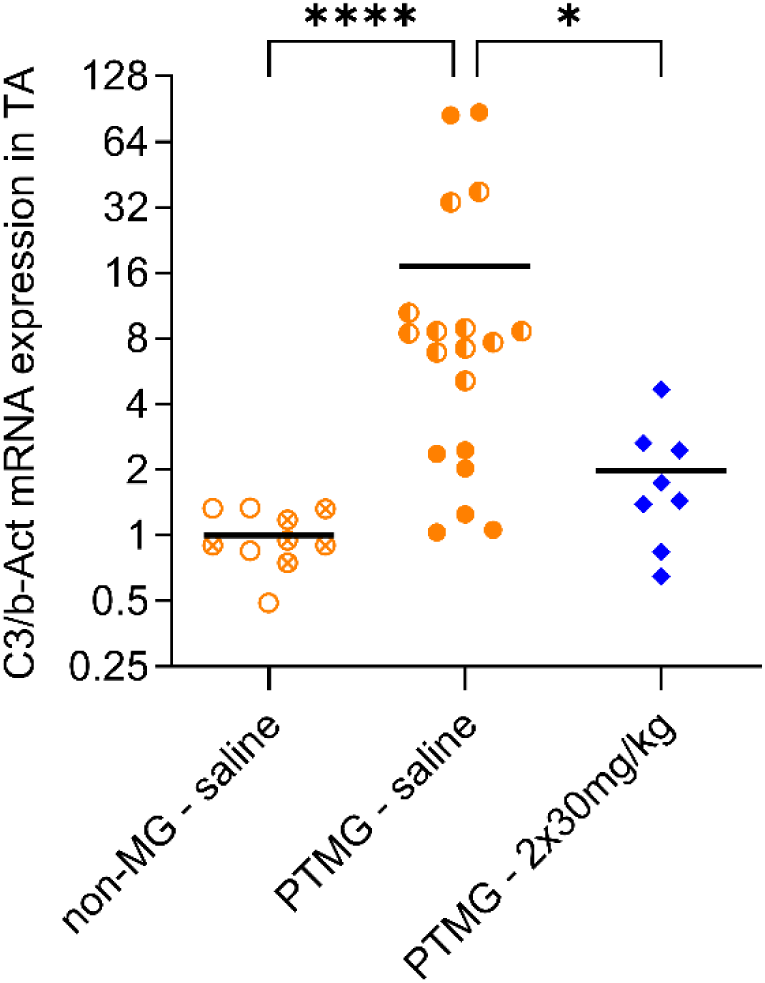
C3 mRNA expression in the TA muscle. RT-qPCR for C3 gene expression was performed with β-Act gene expression as reference in saline treated non-MG (N=10) and PTMG (N=20) animals as well as PTMG animals treated with a double dose of 30 mg/kg C3-siRNA (N=8). All groups were normalized with the non-MG group. Circles with crosses and half-filled circles indicate the animals of the PTMG-2 experiment. Statistical analysis was performed using a Kruskal-Wallis test with post-hoc Dunn’s multiple comparison test; α=0.05, * p=0.0445, **** p<0.0001.

**Figure 8:**
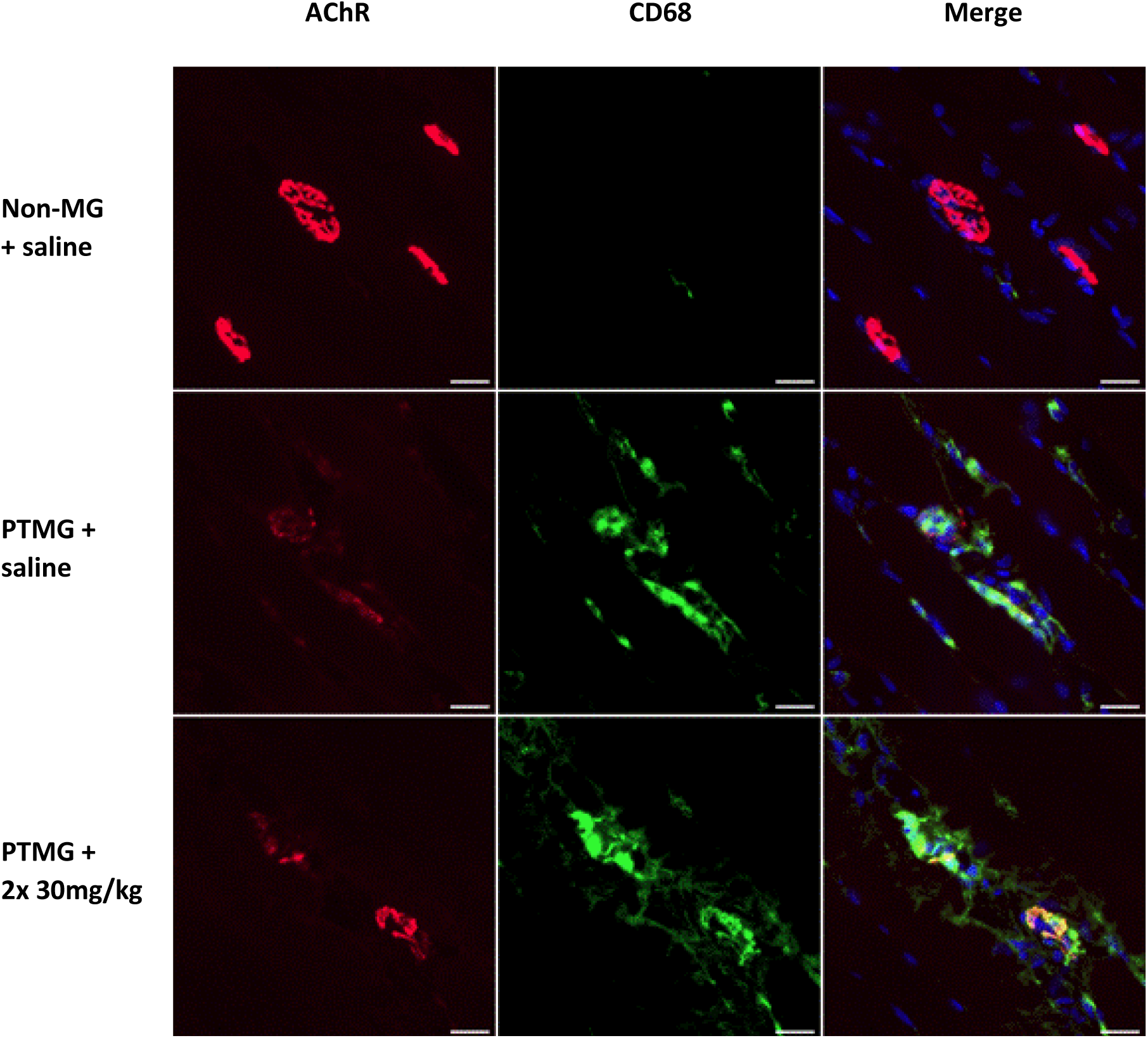
Fluorescence Microscopy Images of Representative Endplates with macrophage infiltration. Representative NMJ images from the non-MG-saline, PTMG-saline, and PTMG 2x 30mg/kg treated groups. Muscle sections were stained for Dapi (blue), AChR (red) and the macrophage marker CD68 (green). Scale bar = 25 µm.

## Discussion

In this study, we investigated the therapeutic effect of a subcutaneously dosed, liver-targeted siRNA against complement factor C3 in the PTMG rat model to assess the role of complement activation in the acute phase of MG. Various treatment paradigms with C3-siRNA were tested: 1x 10 mg/kg, 1x 30 mg/kg, and 2x 30 mg/kg. We observed a clear dose-dependent disease modulation effect of C3-siRNA, when administered prior to PTMG induction. While a single 10 mg/kg dose had minimal impact on disease severity compared to saline-treated PTMG rats, the 2x 30 mg/kg dose paradigm consistently showed a clear protective effect. *In vivo* outcome measurements, including weight loss, grip strength, and disease score, were comparable to those of non-MG animals. These results align with previous findings on C5 inhibition in PTMG rats. Studies using anti-C5 antibodies (21), peptides (24) or siRNAs (27,38) targeting C5 have shown dose-dependent protective effects *in vivo*, including improved grip strength, minimal to no weight loss, and reduced disease score. Similar to the results here, C5 inhibitors reduced MAC deposition at the NMJ, but did not offer full protection (27,38), even though circulating C5 was reduced by up to 95% in one study (38). Similar to the effects observed with C3-siRNA, silencing C5 partially preserved AChR content at the NMJ (27,38) but not to baseline AChR levels seen in non-MG animals (38). These observations raise two key questions:

1. How do PTMG animals, with effective and robust reduction of plasma C3, maintain remarkable clinical stability, including preserved strength and weight, despite complement activity persisting at the NMJ?
2. If systemic C3 is nearly completely depleted, is macrophage infiltration in local tissue responsible for complement production playing a role in MG pathophysiology?

Addressing the first question is relatively straightforward since NMJ functionality remains intact despite a 60% reduction in post-synaptic AChR content before observable clinical signs appear (54). Consequently, neither the internalization of AChR through antigenic modulation, nor the structural degradation of the NMJ caused by complement activation, leads to an immediate loss of bodyweight and strength. As a result, neuromuscular complement deposits can accumulate before translating to clinical signs.

The second question, however, is more complex. As we observed a near complete depletion of circulating C3 protein levels in plasma, it is likely that a local production of complement serves as source for MAC formation. In other studies, rodent and human monocytes and macrophages have been identified as local producers of C3 (56–58) with increased intracellular C3 mRNA levels (58). Macrophage infiltration during the acute phase of MG is well documented (59), and was also demonstrated in our study in PTMG animals, treated with C3-siRNA or saline. Therefore, we could assume that muscle infiltrating macrophages may be responsible for complement production despite effective hepatic C3 suppression. The significant increase of C3 mRNA expression in the muscle observed in PTMG compared to non-MG animals supports this hypothesis. Moreover, macrophage infiltration near the NMJs was observed in PTMG animals treated with C3-siRNA, assumingly primarily activated directly by mAb35 and secondarily due to tissue damage. Although the C3-siRNA treatment substantially prevented the elevation of muscle-located C3 expression in C3-siRNA treated animals, the C3 expression was slightly higher compared to non-MG animals, attributable to macrophage infiltration.

In addition, a C3 bypassing mechanism in complement activation may be important. While C3 production is suppressed with C3-siRNA, hepatic and local cellular C5 production in disease tissue (57) remain unaffected and could be activated directly by the C3 convertase C4bC2a, as it has been observed in a C3 knockout model where MAC-related hemolytic activity was proven. Despite effective C3 knockout, the C3 convertase directly activated C5 with subsequent MAC formation (61).

Although complement deposition was not fully reduced with C3-siRNA treatment, a significant decrease in MAC deposition of 31% compared to saline treated PTMG animals was observed. The substantial improvement in muscle strength and prevention of general disease signs in animals receiving the double dose of C3-siRNA further supports the concept of complement inhibition at the C3 level. In contrast to targeting more downstream complement factors, inhibition of complement at the level of C3 has the potential to reduce the production of the anaphylatoxins, C3a and C5a, which may contribute to a stronger pro-inflammatory profile, with secretion of cytokines including IL-6, IFN-γ, and IFN-α (22,41). Cytokine levels have been shown to be elevated in MG, particularly during MG crises (62,63). Additionally, IL-6 correlates with the activation of lymphocytes and macrophages (63). In the PTMG and the early phase of the EAMG rodent model, macrophage infiltration into muscle tissue is associated with muscle weakness (64,65). Furthermore, macrophage infiltration has been identified in muscle biopsies from MG patients (65), suggesting that local complement activation in MG may also contribute to macrophage infiltration (42) during MG crises.

To advance the experimental setup towards a model more representative of human MG, a follow-up EAMG study with weekly treatment administrations of 30 mg/kg C3-siRNA is strongly recommended. While the PTMG model provides insights into the role of complement during the acute phase of MG, the EAMG model enables the study of the chronic phase of MG. Induced by active immunization of the animals with AChR extracted from the electric ray *Torpedo californica*, the EAMG model mimics the full human disease progression, including the lymphocytic response, complement activation, ultrastructural changes of the NMJ, and progressive muscle weakness, as also seen in patients. To further elucidate the role of complement in MG development and to assess the potential of complement inhibition as a treatment strategy, we propose incorporating both a preventive treatment approach and a post-immunization treatment approach and comparing the therapeutic effect with untreated EAMG animals. To assess the importance of macrophage activation as pathogenic driver during the MG crisis, especially the preventive treatment approach might be of importance, to evaluate the pathogenic potency of local complement production while hepatic C3 is downregulated.

## Conclusion

We demonstrated that suppression of hepatic C3 expression can reduce PTMG-related symptoms in rats. The rapid-acting effect of circulating C3 depletion offers a promising strategy for managing MG crises effectively. These results encourage further investigations of C3 targeting therapies in other MG models such as the chronic EAMG model as a more representative model of the chronic disorder in humans.

## Supplementary data

**Supplementary Figure S1.**
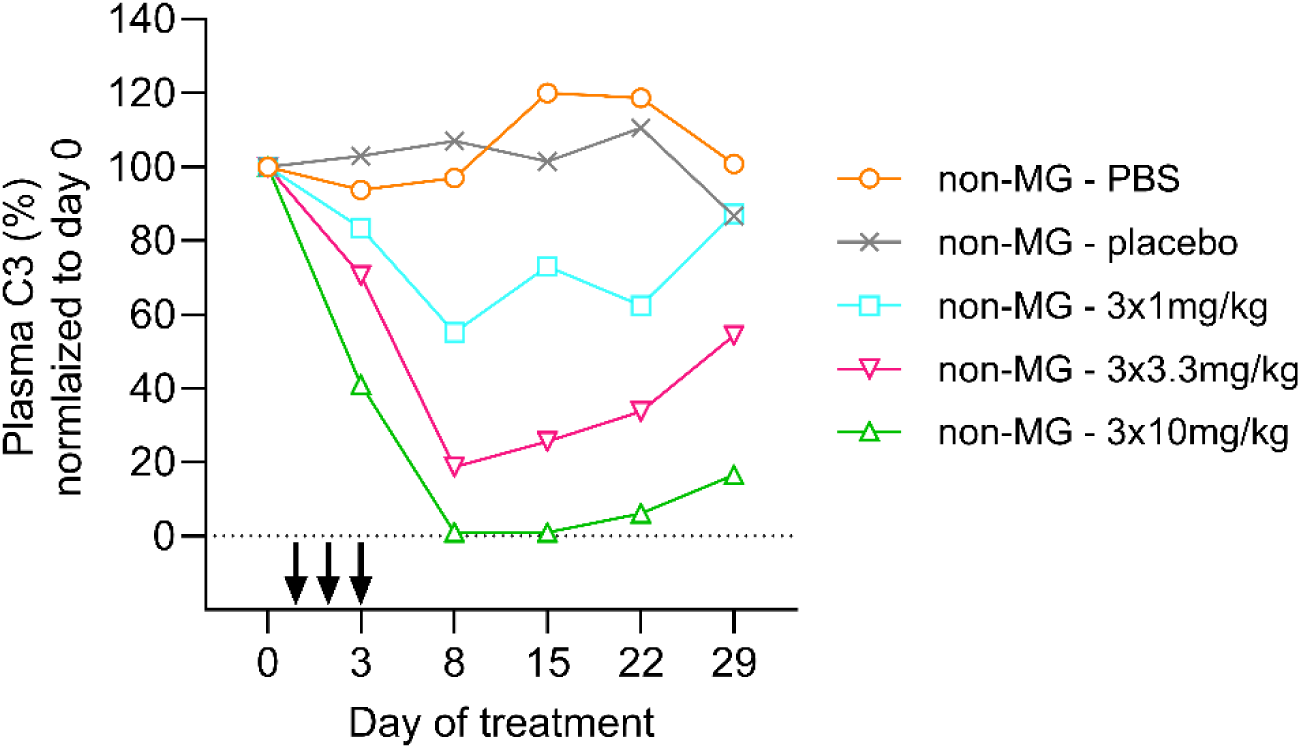
**C3 protein levels in plasma** of healthy animals receiving three consecutive treatment injections, with a cumulative dose equal to a single injection as shown in Fig 1. After baseline measurements on day 0, C3-siRNA was administered on day 1, 2, and 3 (black arrows) and plasma C3 protein levels analyzed on day 3, 8, 15, 22 and 29. Per group N=3. Group averages are shown.

### C3-siRNA detection in liver

Liver samples were collected after euthanasia of each animal, incubated in 2ml cold RNA Stabilization Solution (RNAlater) (ThermoFisher #AM7020) for 24h at 4 °C and subsequently stored at -80 °C upon processing.

To determine the C3-siRNA values in the liver, defrosted liver samples were grinded in 10 µl 0.25% Triton X-100 per 1 mg of tissue at 95 °C, followed by a 10 min incubation at 95 °C including two vortex rounds. The samples were microfuged at highest speed for 20 min at 4 °C, supernatant stored at -20 °C. Reverse Transcription (RT) was performed using the TaqMan MicroRNA Reverse Transcription Kit (Applied Biosystems #4366596) following the manufacturer instructions with the following adaptions. In brief, the single reaction master mix was prepared at 4 °C including 1.5 µl 10X RT Buffer, 3.0 µl 5X RT Primer Pool, 0.15 µl nucleoside trisphosphates (NTPs), 0.2 µl RNase Inhibitor, 1 µl MultiScribe Reverse Transcriptase, 4.15 µl nuclease-free water (all provided with the kit). The lysates were heated to 95 °C for 10 min prior adding 5 µl (0.2 – 2 ng/µl) of lysate to 10 µl master mix. cDNA was produced running the RT program for 30 min at 16 °C, 5 min at 85 °C, and kept subsequently at 4 °C. For the subsequent PCR analysis, the single reaction master mix of 10 µl TaqMan Universal Master Mix II, no UNG (Applied Biosystems #4440049), 0.33 µl Custom TaqMan Small RNA Assay (60X) (Applied Biosystems #4398989), and 7.67 µl nuclease-free water was prepared, vortexed, and 2 µl cDNA sample added. The RT qPCR was initiated with 10 min polymerase activation at 95 °C, followed by 40 cycles of 15 sec at 95 °C and 60 sec at 60 °C. The results were analysed with the delta-delta Ct method.

**Supplementary Figure S2.**
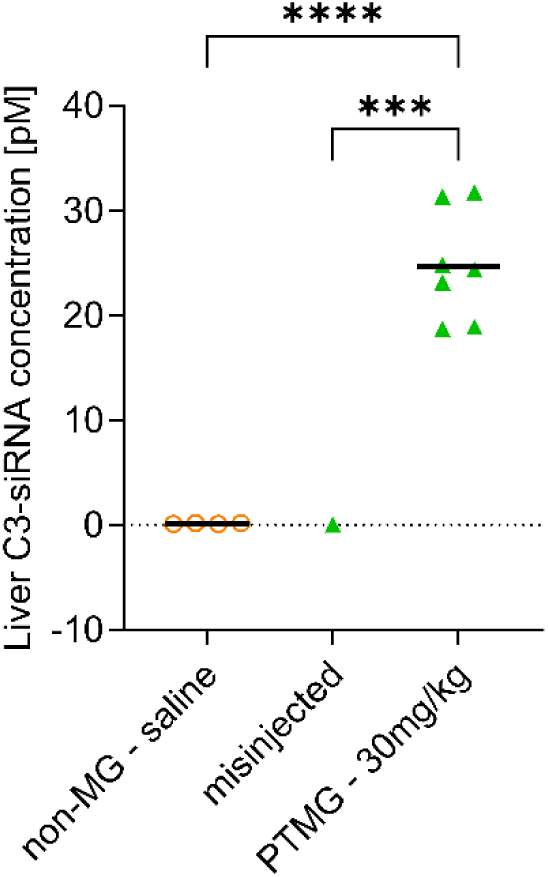
C3-siRNA concentration in the liver after euthanasia. One PTMG animal treated with 30 mg/kg C3-siRNA showed no siRNA in the liver. Assuming a misinjection of C3-siRNA, this animal was excluded from the analysis. Each dot indicates one animal. Statistical analysis was performed using One-Way ANOVA with post-hoc Bonferroni’s multiple comparison test; α=0.05; ***p=0.0009, ****p<0.0001.

**Supplementary Figure S3.**
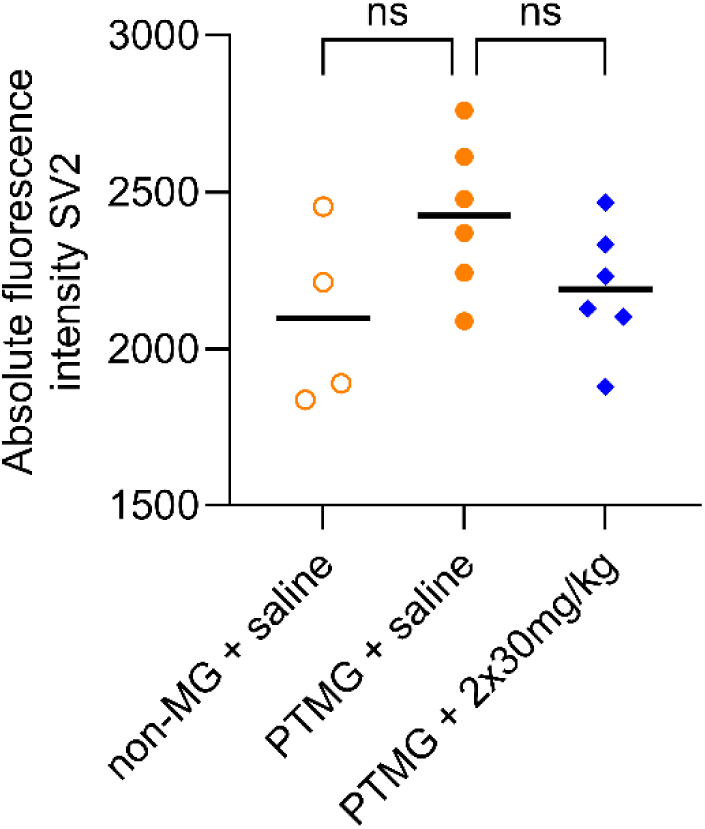
Quantitative immunofluorescence data for SV2. The presynaptic SV2 remains unaffected by PTMG, as indicated by similar absolute immunofluorescence intensities across all treatment groups (PTMG and non-MG), whether they received saline or a double injection of 30 mg/kg C3siRNA. N=4 non-MG animals, N=6 PTMG-saline animals, N=6 PTMG-2x 30 mg/kg animals. Each dot represents one animal. One-Way ANOVA with post hoc Bonferroni’s multiple comparison test; α=0.05.

### Clinical chemistry

Clinical chemistry was analyzed in healthy rats by Apellis Pharmaceuticals in the laboratories of Wuxi AppTech, US. Five healthy, female Lewis rats of 10 weeks of age were obtained from Hilltop Lab Animals, Inc., US, and sc injected with 30 mg/kg C3-siRNA weekly over a period of 6 weeks. Hepatotoxicity data in plasma for aspartate transferase (AST), alkaline phosphatase (ALP), alanine transaminase (ALT) and glutamate dehydrogenase (GLDH) were obtained before any treatment injection on day 0, day 22, and day 43. Data of non-injected, naïve animals (N=6) were obtained only on day 43. The blood was collected from the jugular vein or through cardiac puncture (day 43) into K2-EDTA tubes., centrifuged for 5 min with 3000 g at 4 °C.

**Supplementary Figure S4.**
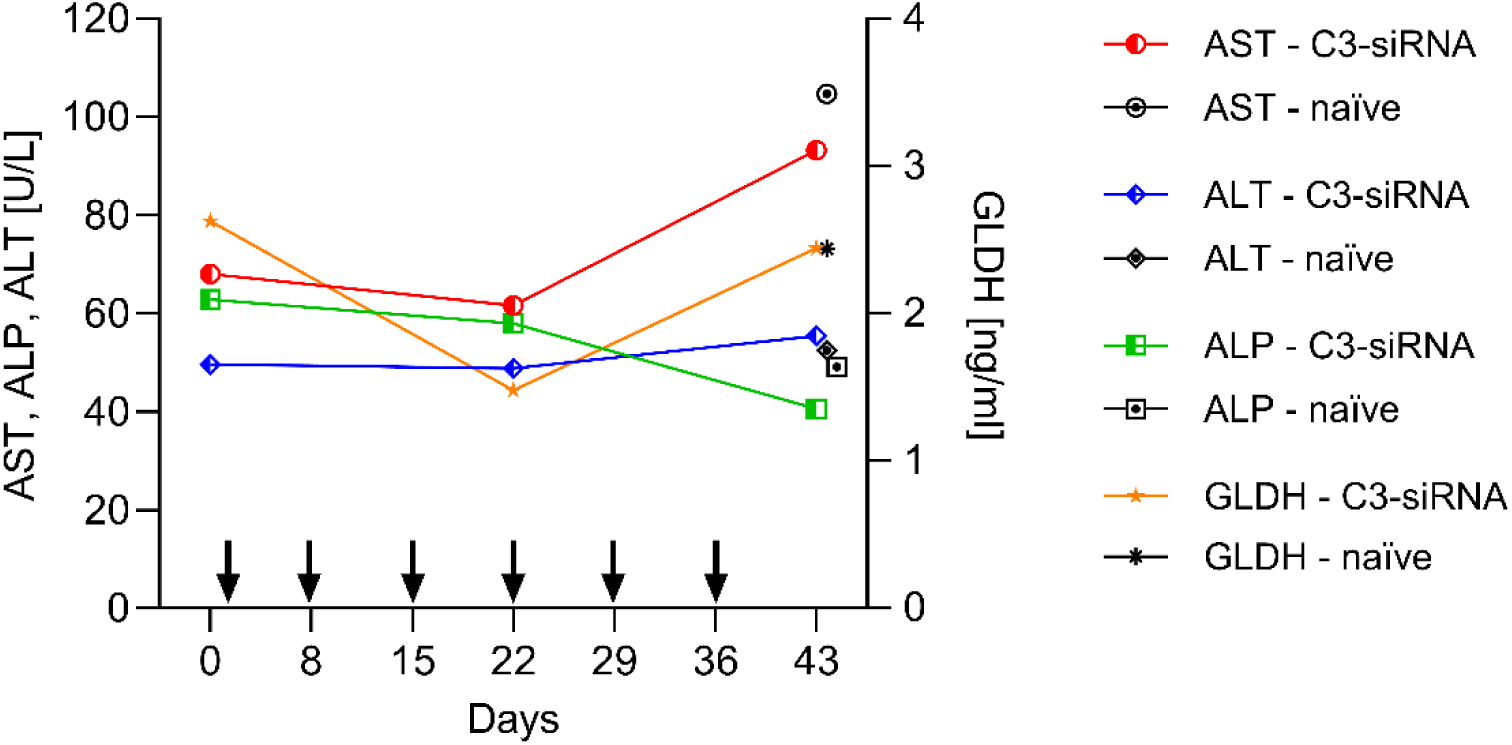
Plasma liver enzyme levels to assess hepatotoxicity of the C3-siRNA. Aspartate transferase (AST), alkaline phosphatase (ALP), alanine transaminase (ALT) and glutamate dehydrogenase (GLDH) were analyzed in plasma on day 0, day 22 and day 43. Data of untreated, naïve rats were used as controls on day 43. C3-siRNA treatment was administered on day 1, 8, 15, 22, 29 and 36. N=5 healthy 30 mg/kg C3-siRNA treated animals (coloured), N=6 untreated, naïve animals (black). Each dot represents the group’s mean. Unpaired t-test; α=0.05.

No significant changes in the liver enzyme levels AST, ALP, ALT or GLDH were observed in C3-siRNA treated rats between day 0 and D43. Further, no significant difference of the enzyme levels between C3-siRNA treated and naïve rats on day 43 was detected. Additionally to the liver enzymes, parameters of general toxicology to detect metabolic irregularities were analyzed at several timepoints of this experiment (data not shown). None of these parameters indicated toxicity of the C3-siRNA.

### C3 mRNA expression in the rat liver after treatment with C3-siRNA

Hepatic gene expression of C3 after C3-siRNA treatment was assessed in context of the dose range study described in Table 1 and (Supplementary) Fig. 1. Liver C3 mRNA levels were determined at two timepoints after treatment injection: 3 days after treatment administration, including only the single-injected animals (1x3 / 1x10 / 1x3 mg/kg C3-siRNA, and PBS; per group N=3); and 30 days after treatment injection, including all animal groups with single- and triple-injected animals (1x3 / 1x10 / 1x3 mg/kg, 3x1 / 3x3.3 / 3x10 mg/kg C3-siRNA, PBS, and non-targeting; per group N=3). The analyses were performed by Axolabs GmbH, Kulmbach, Germany. The results were analysed with the delta-delta Ct method.

**Supplementary Figure S5.**
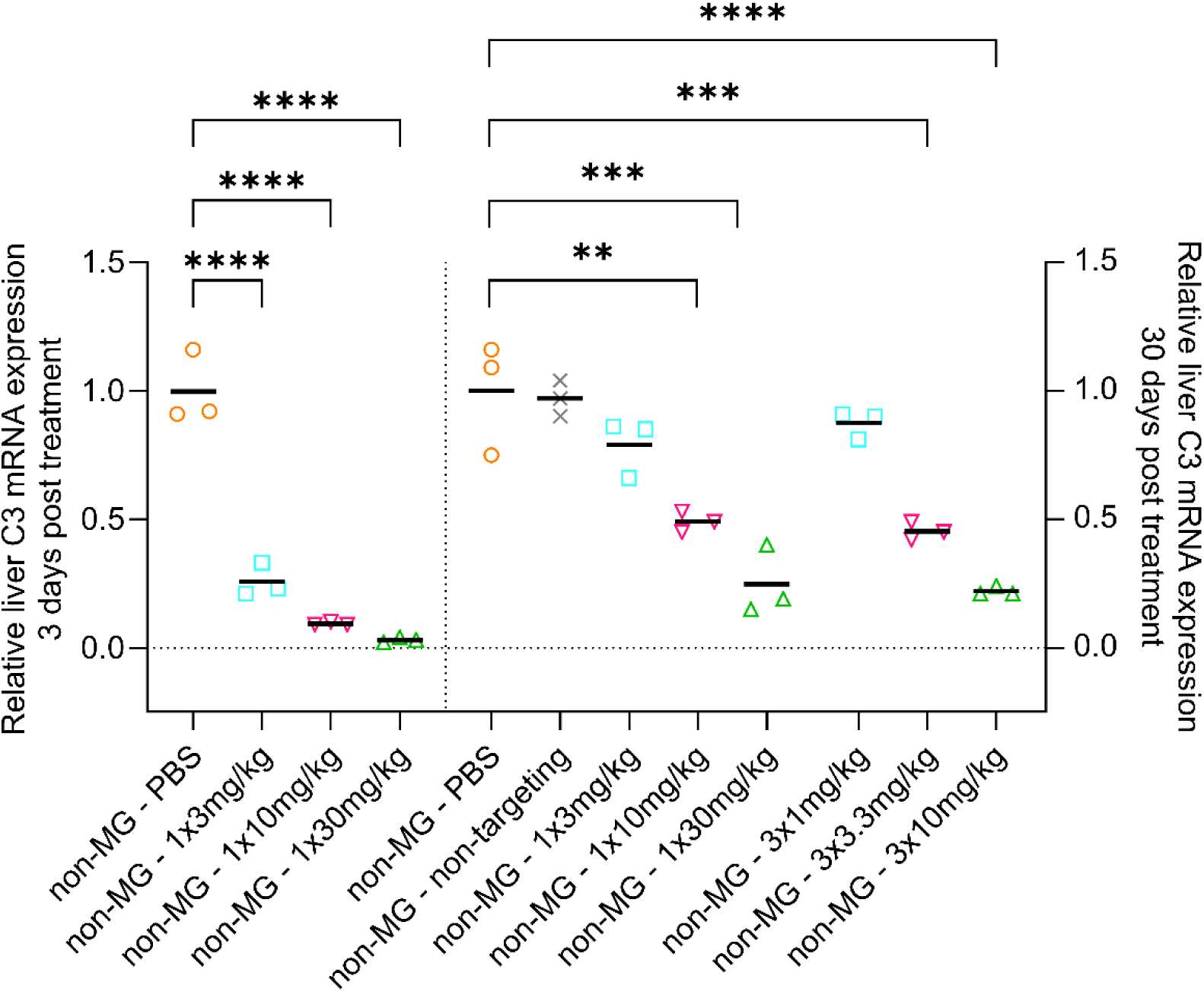
Hepatic C3 mRNA expression 3 and 30 days after C3-siRNA injection. Three days after treatment injection, hepatic C3 mRNA expression was significantly repressed in all C3-siRNA treated animals (left). 30 days after C3-siRNA injection, the expression was still reduced to 50% and ca. 25% in animals that received 10 or 30 mg/kg C3-siRNA (single and cumulative) compared to control animals. Each dot indicates one animal. Statistical analysis as performed using One-Way ANOVA with post-hoc Bonferroni’s multiple comparison test; α=0.05; **p=0.003, ***p<0.0004, ****p<0.0001.

In animals receiving a single injection of C3-siRNA, C3 mRNA expression was effectively reduced three days after treatment injection with a visible dose-dependent effect (Supplementary Fig. 5). The highest dose of 30 mg/kg C3-siRNA inhibited C3 expression to below 5% compared to the expression in PBS-treated animals (p<0.0001). 30 days after treatment injection, the C3 expression was still impaired and reduced to 50% and 25% in animals treated with 10 mg/kg or 30 mg/kg C3-siRNA, respectively. Those animals receiving triple-injections expressed similar levels of C3 mRNA on day 30 as those receiving the same dose in a single injection. From these results it can be concluded that the C3-siRNA was successfully transported to the liver and effectively suppresses hepatic C3 production.

## Acknowledgements

We thank Barbie Machiels for technical assistance with the RIAs; further we thank Veerle Horsting for performing the immunofluorescent stainings.

## Funding

This study was supported by Apellis Pharmaceuticals, Inc., Waltham, Massachusetts, USA.

## Conflict of interest

E. Richardson, T. Barbour and David Eyerman are current employees of Apellis Pharmaceuticals. Lukas Scheibler is a former employee of Apellis Pharmaceuticals. M.H. De Baets declares to be a consultant for Apellis Pharmaceuticals.

## References

1. Verschuuren JJ, Huijbers MG, Plomp JJ, Niks EH, Molenaar PC, Martinez-Martinez P, et al. Pathophysiology of myasthenia gravis with antibodies to the acetylcholine receptor, muscle-specific kinase and low-density lipoprotein receptor-related protein 4. Autoimmun Rev. 2013/03/29. 2013;12(9):918–23.

2. Gilhus NE, Skeie GO, Romi F, Lazaridis K, Zisimopoulou P, Tzartos S. Myasthenia gravis — autoantibody characteristics and their implications for therapy. Nat Rev Neurol 2016 125. 2016 Apr 22;12(5):259–68.

3. Lazaridis K, Tzartos SJ. Myasthenia Gravis: Autoantibody Specificities and Their Role in MG Management. Vol. 11, Frontiers in Neurology. Frontiers Media S.A.; 2020.

4. Rødgaard A, Nielsen FC, Djurup R, Somnier F, Gammeltoft S. Acetylcholine receptor antibody in myasthenia gravis: predominance of IgG subclasses 1 and 3. Clin Exp Immunol. 1987 Jan;67(1):82.

5. Losen M, Martínez-Martínez P, Phernambucq M, Schuurman J, Parren PWHI, De Baets MH. Treatment of myasthenia gravis by preventing acetylcholine receptor modulation. In: Annals of the New York Academy of Sciences. Blackwell Publishing Inc.; 2008. p. 174–9.

6. Romi F, Kristoffersen EK, Aarli JA, Gilhus NE. The role of complement in myasthenia gravis: Serological evidence of complement consumption in vivo. J Neuroimmunol. 2005;158(1– 2):191–4.

7. Tüzün E, Christadoss P. Complement associated pathogenic mechanisms in myasthenia gravis. Autoimmun Rev. 2013;12:904–11.

8. Howard Jr. JF. Myasthenia gravis: the role of complement at the neuromuscular junction. Ann N Y Acad Sci. 2017/12/22. 2018;1412(1):113–28.

9. Martinez-Martinez P, Phernambucq M, Steinbusch L, Schaeffer L, Berrih-Aknin S, Duimel H, et al. Silencing rapsyn in vivo decreases acetylcholine receptors and augments sodium channels and secondary postsynaptic membrane folding. Neurobiol Dis. 2009;35(1):14–23.

10. Losen M, Stassen MH, Martinez-Martinez P, Machiels BM, Duimel H, Frederik P, et al. Increased expression of rapsyn in muscles prevents acetylcholine receptor loss in experimental autoimmune myasthenia gravis. Brain. 2005;128(Pt 10):2327–37.

11. Martinez-Martinez P, Losen M, Duimel H, Frederik P, Spaans F, Molenaar P, et al. Overexpression of rapsyn in rat muscle increases acetylcholine receptor levels in chronic experimental autoimmune myasthenia gravis. Am J Pathol. 2007/01/27. 2007;170(2):644–57.

12. Gomez AM, Stevens JA, Mane-Damas M, Molenaar P, Duimel H, Verheyen F, et al. Silencing of Dok-7 in Adult Rat Muscle Increases Susceptibility to Passive Transfer Myasthenia Gravis. Am J Pathol. 2016/09/24. 2016;186(10):2559–68.

13. Engel AG, Sahasi K, Lindstrom JM, Lambert EH, Lennon VA SH, Engel AG, Sakakibara H, Sahashi K, Lindstrom JM, Lambert EH, et al. Passively transferred experimental autoimmune myasthenia gravis. equential and quantitative study of the motor end-plate fine structure and ultrastructural localization of immune complexes (IgG and C3), and of the acetylcholine receptor. Neurology. 1979;29(2):10.

14. Sahashi K, Engel AG, Lambert EH, Howard FM. Ultrastructural localization of the terminal and lytic ninth complement component (C9) at the motor end-plate in myasthenia gravis. J Neuropathol Exp Neurol. 1980;39(2):160–72.

15. Mané-Damas M, Molenaar PC, Ulrichts P, Marcuse F, De Baets MH, Martinez-Martinez P, et al. Novel treatment strategies for acetylcholine receptor antibody-positive myasthenia gravis and related disorders. Autoimmun Rev. 2022 Jul 1;21(7).

16. Maggi L, Mantegazza R. Treatment of myasthenia gravis: Focus on pyridostigmine. Clin Drug Investig. 2011 Aug 24;31(10):691–701.

17. Sanders DB, Wolfe GI, Benatar M, Evoli A, Gilhus NE, Illa I, et al. International consensus guidance for management of myasthenia gravis: Executive summary. Neurology. 2016 Jul 7;87(4):419.

18. Schneider-Gold C, Hagenacker T, Melzer N, Ruck T. Understanding the burden of refractory myasthenia gravis. Ther Adv Neurol Disord. 2019 Feb 1;12.

19. Wolfe GI, Kaminski HJ, Aban IB, Minisman G, Kuo H-C, Marx A, et al. Randomized Trial of Thymectomy in Myasthenia Gravis. N Engl J Med. 2016 Aug 11;375(6):511–22.

20. Albazli K, Kaminski HJ, Howard JF, Howard Jr. JF. Complement Inhibitor Therapy for Myasthenia Gravis. Front Immunol. 2020/06/26. 2020 Jun 3;11:917.

21. Zhou Y, Gong B, Lin F, Rother RP, Medof ME, Kaminski HJ. Anti-C5 antibody treatment ameliorates weakness in experimentally acquired myasthenia gravis. J Immunol. 2007/12/07. 2007;179(12):8562–7.

22. Tüzün E, Scott BG, Goluszko E, Higgs S, Christadoss P. Genetic Evidence for Involvement of Classical Complement Pathway in Induction of Experimental Autoimmune Myasthenia Gravis. J Immunol. 2003 Oct 1;171(7):3847–54.

23. Christadoss P. C5 gene influences the development of murine myasthenia gravis. J Immunol. 1988/04/15. 1988;140(8):2589–92.

24. Soltys J, Kusner LL, Young A, Richmonds C, Hatala D, Gong B, et al. Novel complement inhibitor limits severity of experimentally myasthenia gravis. Ann Neurol. 2009 Jan;65(1):67– 75.

25. Huda R, Tüzün E, Christadoss P. Complement C2 siRNA mediated therapy of myasthenia gravis in mice. J Autoimmun. 2013;42:94–104.

26. Biesecker G, Gomez CM. Inhibition of acute passive transfer experimental autoimmune myasthenia gravis with Fab antibody to complement C6. J Immunol. 1989/04/15. 1989;142(8):2654–9.

27. Kusner LL, Yucius K, Sengupta M, Sprague AG, Desai D, Nguyen T, et al. Investigational RNAi Therapeutic Targeting C5 Is Efficacious in Pre-clinical Models of Myasthenia Gravis. Mol Ther - Methods Clin Dev. 2019 Jun 14;13:484–92.

28. Tüzün E, Huda R, Christadoss P. Complement and cytokine based therapeutic strategies in myasthenia gravis. Vol. 37, Journal of Autoimmunity. Academic Press; 2011. p. 136–43.

29. Narayanaswami P, Sanders DB, Wolfe G, Benatar M, Cea G, Evoli A, et al. International Consensus Guidance for Management of Myasthenia Gravis: 2020 Update. Neurology. 2021 Jan 19;96(3):114–22.

30. Rystiggo | European Medicines Agency (EMA) [Internet]. [cited 2024 Nov 4]. Available from: https://www.ema.europa.eu/en/medicines/human/EPAR/rystiggo

31. Zilbrysq | European Medicines Agency (EMA) [Internet]. [cited 2024 Nov 4]. Available from: https://www.ema.europa.eu/en/medicines/human/EPAR/zilbrysq

32. Kusner LL, Losen M, Vincent A, Lindstrom J, Tzartos S, Lazaridis K, et al. Guidelines for pre-clinical assessment of the acetylcholine receptor--specific passive transfer myasthenia gravis model-Recommendations for methods and experimental designs. Exp Neurol. 2015/03/10. 2015;270:3–10.

33. Losen M, Martinez-Martinez P, Molenaar PC, Lazaridis K, Tzartos S, Brenner T, et al. Standardization of the experimental autoimmune myasthenia gravis (EAMG) model by immunization of rats with Torpedo californica acetylcholine receptors - Recommendations for methods and experimental designs. Exp Neurol. 2015 Jan 2;270:18–28.

34. Tuzun E, Berrih-Aknin S, Brenner T, Kusner LL, Le Panse R, Yang H, et al. Guidelines for standard preclinical experiments in the mouse model of myasthenia gravis induced by acetylcholine receptor immunization. Exp Neurol. 2015 Aug 1;270:11–7.

35. Tüzün E, Li J, Saini SS, Yang H, Christadoss P. Pros and cons of treating murine myasthenia gravis with anti-C1q antibody. J Neuroimmunol. 2007 Jan 1;182(1–2):167–76.

36. Christadoss P, Tüzün E, Li J, Saini SS, Yang H. Classical complement pathway in experimental autoimmune myasthenia gravis pathogenesis. Ann N Y Acad Sci. 2008;1132:210–9.

37. Kusner LL, Satija N, Cheng G, Kaminski HJ. Targeting therapy to the neuromuscular junction: Proof of concept. Muscle Nerve. 2014 May 1;49(5):749–56.

38. Kuboi Y, Suzuki Y, Motoi S, Matsui C, Toritsuka N, Nakatani T, et al. Identification of potent siRNA targeting complement C5 and its robust activity in pre-clinical models of myasthenia gravis and collagen-induced arthritis. Mol Ther Nucleic Acids. 2023 Mar 14;31:339–51.

39. Study Details | NCT05070858 | A Study to Test How Safe Pozelimab and Cemdisiran Combination Therapy and Cemdisiran Alone Are and How Well They Work in Adult Patients With Generalized Myasthenia Gravis | ClinicalTrials.gov [Internet]. [cited 2025 Aug 31]. Available from: https://clinicaltrials.gov/study/NCT05070858?intr=Cemdisiran&rank=2

40. Tüzün E, Huda R, Christadoss P. Complement and cytokine based therapeutic strategies in myasthenia gravis. J Autoimmun. 2011;37(2):136–43.

41. Deng C, Goluszko E, Tüzün E, Yang H, Christadoss P. Resistance to Experimental Autoimmune Myasthenia Gravis in IL-6-Deficient Mice Is Associated with Reduced Germinal Center Formation and C3 Production. J Immunol. 2002 Jul 15;169(2):1077–83.

42. Zarantonello A, Revel M, Grunenwald A, Roumenina LT. C3-dependent effector functions of complement. Immunol Rev. 2023 Jan 1;313(1):120.

43. Ricklin D, Reis ES, Mastellos DC, Gros P, Lambris JD. Complement component C3 - The “Swiss Army Knife” of innate immunity and host defense. Immunol Rev. 2016 Nov 1;274(1):33.

44. Xu L, Xu H, Chen S, Jiang W, Afridi SK, Wang Y, et al. Inhibition of complement C3 signaling ameliorates locomotor and visual dysfunction in autoimmune inflammatory diseases. Mol Ther. 2023 Sep 9;31(9):2715.

45. Soltys J, Wu X. Complement regulatory protein Crry deficiency contributes to the antigen specific recall response in experimental autoimmune myasthenia gravis. J Inflamm (Lond). 2012;9(1).

46. Setten RL, Rossi JJ, Han S ping. The current state and future directions of RNAi-based therapeutics. Nat Rev Drug Discov 2019 186. 2019 Mar 7;18(6):421–46.

47. Fitzgerald K, White S, Borodovsky A, Bettencourt BR, Strahs A, Clausen V, et al. A Highly Durable RNAi Therapeutic Inhibitor of PCSK9. N Engl J Med. 2017 Jan 5;376(1):41–51.

48. Nishimura J, Yamamoto M, Hayashi S, Ohyashiki K, Ando K, Brodsky AL, et al. Genetic variants in C5 and poor response to eculizumab. N Engl J Med. 2014 Feb 13;370(7):632–9.

49. Ranasinghe P, Addison ML, Dear JW, Webb DJ. Small interfering RNA: Discovery, pharmacology and clinical development—An introductory review. Br J Pharmacol. 2023 Nov 1;180(21):2697–720.

50. Alshaer W, Zureigat H, Al Karaki A, Al-Kadash A, Gharaibeh L, Hatmal MM, et al. siRNA: Mechanism of action, challenges, and therapeutic approaches. Eur J Pharmacol. 2021 Aug 15;905:174178.

51. Springer AD, Dowdy SF. GalNAc-siRNA Conjugates: Leading the Way for Delivery of RNAi Therapeutics. Nucleic Acid Ther. 2018;28(3):109–18.

52. Matsuda S, Keiser K, Nair JK, Charisse K, Manoharan RM, Kretschmer P, et al. siRNA Conjugates Carrying Sequentially Assembled Trivalent N-Acetylgalactosamine Linked Through Nucleosides Elicit Robust Gene Silencing In Vivo in Hepatocytes. ACS Chem Biol. 2015;10(5):1181–7.

53. Zhang L, Liang Y, Liang G, Tian Z, Zhang Y, Liu Z, et al. The therapeutic prospects of N-acetylgalactosamine-siRNA conjugates. Front Pharmacol. 2022;13(December):1–13.

54. Losen M, Martinez-Martinez P, Molenaar PC, Lazaridis K, Tzartos S, Brenner T, et al. Standardization of the experimental autoimmune myasthenia gravis (EAMG) model by immunization of rats with Torpedo californica acetylcholine receptors--Recommendations for methods and experimental designs. Exp Neurol. 2015/03/23. 2015;270:18–28.

55. Tse N, Morsch M, Ghazanfari N, Cole L, Visvanathan A, Leamey C, et al. The Neuromuscular Junction: Measuring Synapse Size, Fragmentation and Changes in Synaptic Protein Density Using Confocal Fluorescence Microscopy. J Vis Exp. 2014 Dec 26;(94):e52220.

56. Kiss MG, Papac-Miličević N, Porsch F, Tsiantoulas D, Hendrikx T, Takaoka M, et al. Cell-autonomous regulation of complement C3 by factor H limits macrophage efferocytosis and exacerbates atherosclerosis. Immunity. 2023 Aug 8;56(8):1809–1824.e10.

57. Niyonzima N, Rahman J, Kunz N, West EE, Freiwald T, Desai J V., et al. Mitochondrial C5aR1 activity in macrophages controls IL-1β production underlying sterile inflammation. Sci Immunol. 2021 Dec 12;6(66):eabf2489.

58. Kolev M, West EE, Kunz N, Chauss D, Moseman EA, Rahman J, et al. Diapedesis-Induced Integrin Signaling via LFA-1 Facilitates Tissue Immunity by Inducing Intrinsic Complement C3 Expression in Immune Cells. Immunity. 2020 Mar 17;52(3):513–527.e8.

59. Huda R. Inflammation and autoimmune myasthenia gravis. Front Immunol. 2023 Jan 30;14:1110499.

60. West EE, Kemper C. Complosome — the intracellular complement system. Nat Rev Nephrol 2023 197. 2023 Apr 13;19(7):426–39.

61. Zhang L, Dai Y, Huang P, Saunders TL, Fox DA, Xu J, et al. Absence of complement component 3 does not prevent classical pathway-mediated hemolysis. Blood Adv. 2019;3(12):1808–14.

62. Huan X, Luo S, Zhong H, Zheng X, Song J, Zhou L, et al. In-depth peripheral CD4+ T profile correlates with myasthenic crisis. Ann Clin Transl Neurol. 2021 Apr 1;8(4):749.

63. Uzawa A, Kuwabara S, Suzuki S, Imai T, Murai H, Ozawa Y, et al. Roles of cytokines and T cells in the pathogenesis of myasthenia gravis. Clin Exp Immunol. 2021 Feb 10;203(3):366–74.

64. Engel AG, Tsujihata M, Lindstrom JM, Lennon VA. The motor end plate in myasthenia gravis and in experimental autoimmune myasthenia gravis. A quantitative ultrastructural study. Ann N Y Acad Sci. 1976;274(1):60–79.

65. Corey AL, Richman DP, Shuman CA, Gomez CM, Arnason BGW. Use of monoclonal antiacetylcholine receptor antibodies to investigate the macrophage inflammation of acute experimental myasthenia gravis: Refractoriness to a second episode of acute disease. Neurology. 1985;35(10):1455–60.

